# Phylogenomic approaches reveal how a climatic inversion and glacial refugia shape patterns of diversity in an African rain forest tree species

**DOI:** 10.1101/807727

**Authors:** Andrew J. Helmstetter, Biowa E. N. Amoussou, Kevin Bethune, Narcisse G. Kandem, Romain Glèlè Kakaï, Bonaventure Sonké, Thomas L. P. Couvreur

## Abstract

The world’s second largest expanse of tropical rain forest is in Central Africa and it harbours enormous species diversity. Population genetic studies have consistently revealed significant structure across central African rain forest plants, in particular a North-South genetic discontinuity close to the equator at the level of a climatic inversion. Here, we take a phylogeographic approach using 351 nuclear markers in 112 individuals across the distribution of the African rain forest tree species *Annickia affinis* (Annonaceae). We show for the first time that the North-South divide is the result of a single major colonisation event across the climatic inversion from an ancestral population located in Gabon. We suggest that differences in ecological niche of populations distributed either side of this inversion may have contributed to this phylogenetic discontinuity. We find evidence for inland dispersal, predominantly in northern areas, and variable demographic histories among genetic clusters, indicating that populations responded differently to past climate change. We show how newly-developed genomic tools can provide invaluable insights into our understanding of tropical rain forest evolutionary dynamics.

## 1 INTRODUCTION

Tropical rain forests (TRFs) possess an incredibly diverse flora and fauna making up half of the world’s biodiversity. Understanding how this diversity is generated is critical if we are to protect it (Mace et al. 2003). Central Africa hosts the world’s second largest continuous extent of TRF (Linder 2001). Climatic fluctuations during the Pleistocene and associated glacial forest refugia are suggested to have influenced how genetic diversity is distributed in Central African rain forests (CAR) (Hardy et al. 2013). However, the nature (Anhuf et al. 2006; Diamond and Hamilton 1980; Maley 1996; Bonnefille 2007) and importance of forest refugia continue to be intensely debated (Cowling et al. 2008; Lezine et al. 2019). Population genetic studies within CAR plant species document differing levels of response to past climatic fluctuations (reviewed in Hardy et al. 2013). Conversely, one major phylogeographic pattern common to many CAR plant species studied is the existence of a phylogeographic barrier along a North-South axis around 0-3°N (see Fig. 1A; Hardy et al. 2013; Faye et al. 2016; Heuertz et al. 2014). There appears to be no visible geographic barrier to explain this break as continuous rain forest exists across the entire area. This North-South phylogeographic barrier corresponds, however, to the central African climatic hinge, an inversion zone between the northern and southern rainy seasons (Hardy et al; 2013). Interestingly, this barrier is rarely recovered in phylogeographic studies of animals and thus seems to affect a greater effect on plants groups (e.g. Fuchs and Bowie 2015; Bohoussou et al. 2015; Bell et al. 2017; Blatrix et al. 2017).

**Fig. 1.**
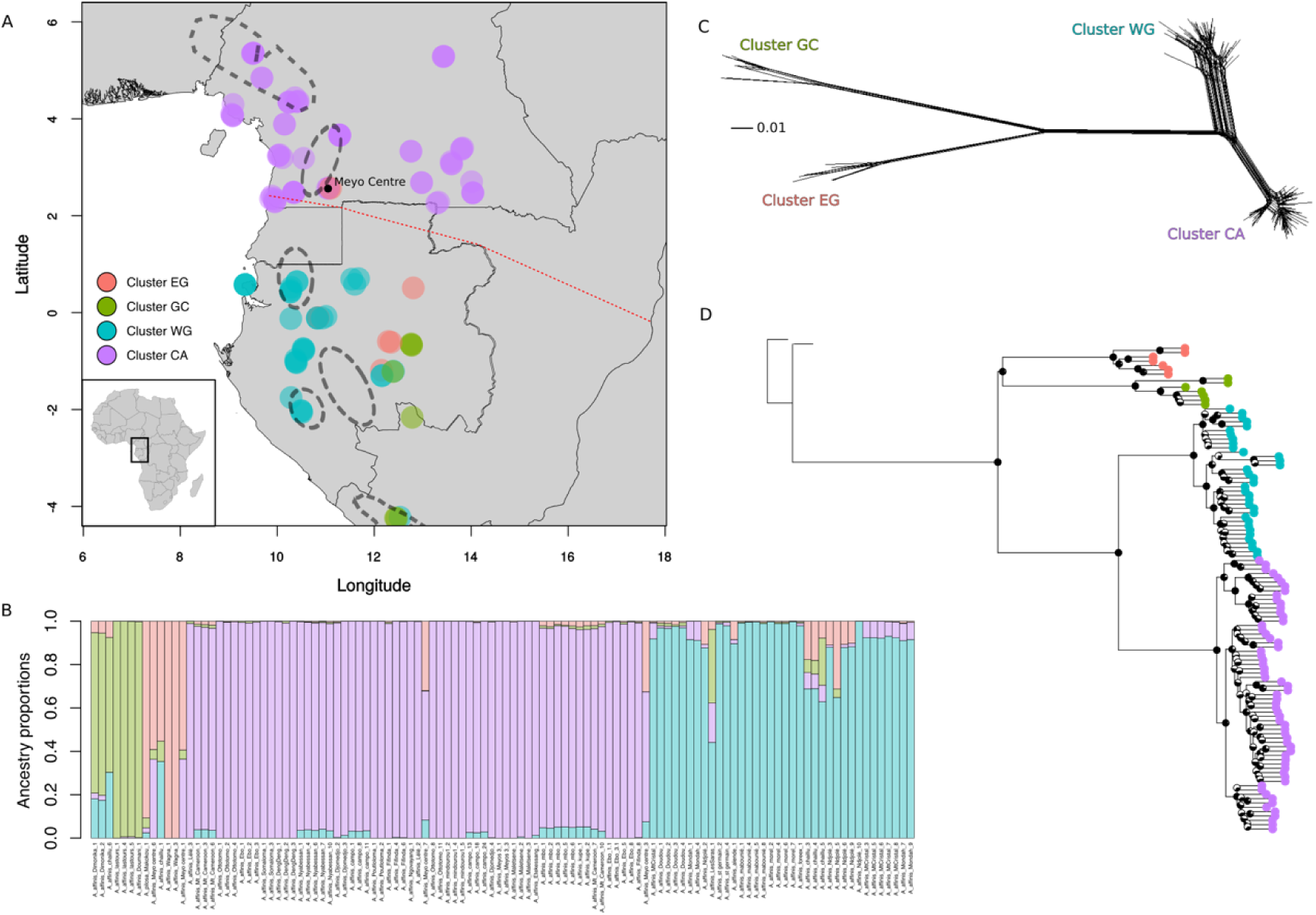
(A) Map of the study region showing genetic clusters inferred using Discriminant analysis of principal components (DAPC, k=4). Individuals are colour coded by cluster membership. Superimposed upon the map are the locations of putative glacial refugia (adapted from Faye et al. 2016). The climatic hinge is shown by a dashed red line. Inset is a map of the African continent showing highlighting the study area. (B) Barplot of ancestry proportions inferred using TESS (k=4). Colours were made to correspond to those in Fig. 1A as clustering was almost identical between approaches. A single individual, “A_affinis_Ndjole_5”, was inferred as cluster EG (red) in DAPC but TESS suggests the majority of its ancestry is instead from cluster WG (blue). Individuals from the “Meyo Centre” show evidence of admixture between clusters (red and purple) across the North-South climatic inversion. (C) Phylogenetic network among *A. affinis* individuals constructed in splitstree using NeighbourNet algorithm based on 6787 SNPs. (D) ASTRAL tree representing relationships among *Annickia affinis* samples, rooted on two *A. polycarpa* samples. Local posterior probabilities are shown as pie charts on nodes and tips are coloured based on genetic clustering results.

Three main hypotheses have been suggested to explain how this North-South genetic discontinuity originated (Hardy et al. 2013). First, TRF might have disappeared (repeatedly) along the climatic hinge during past climatic fluctuations, isolating allopatric north/south populations, which subsequently recolonised the area. In this case we would expect past climate models to show that suitable habitat disappeared along the hinge during glacial periods when forests retract. Second, because seasons are inverted across the hinge, flowering time might be displaced between northern and southern populations preventing interbreeding and gene flow. If flowering occurs at different times of the year either side of the inversion it may indicate this scenario helped shape patterns in genetic structure. Third, successful colonisation across the climatic hinge may be limited by factors such as environmental differences (e.g. changing levels of water stress). If this was the case, we would expect ecological niches of populations either side of the hinge to be significantly different. At present, little is known about the relative importance of these three scenarios so we aim to identify which, if any, have contributed to this common genetic discontinuity.

The phylogeographic approach (Avise et al. 1987) can unravel the history of populations, and ultimately uncover which processes have shaped current patterns of diversity. It is therefore an ideal framework to study how the climatic inversion, and other factors, have shaped patterns of intraspecific diversity in CAR plants. High-throughput sequencing allows the generation of phylogeographic datasets consisting of large numbers of independently segregating nuclear loci that can account for coalescent stochasticity (Edwards et al. 2016) where studies with small numbers of markers fall short. Genomic data can also be used to reconstruct the spatial evolutionary history of species by inferring phylogenetic trees among populations and dispersal dynamics. These approaches can help us to understand relationships among populations on either side of the climatic inversion and how often lineages traversed this barrier.

If glacial refugia in areas proposed by Maley (1996) and Anhuf et al. (2006) have played an important role in CAR plant dynamics we would expect to find evidence of eastward dispersal because most putative CAR refugia are located in the Atlantic Guineo-Congolian region. For example, within the palm species *Podococcus barteri*, modelling past ranges and genetic data supported the hypothesis of one large coastal refugia in western Gabon and southwestern Cameroon (Faye et al. 2016). Furthermore, we would expect population size to increase towards the present as populations spread out from climatically stable areas. If the climatic inversion acts as a barrier, phylogeographic approaches should reveal limited numbers lineages that dispersed across the region from 0-3°N. The nature of this barrier can then be determined by a combination of phenological data and ecological niche modelling approaches as detailed above.

To develop upon our current understanding of the phylogeographic patterns introduced above, we present, for the first time using nuclear phylogenomic approaches, the evolutionary dynamics of a central African tree species, *Annickia affinis.* This species belongs to the pantropical plant family Annonaceae (Chatrou et al. 2012) growing up to 30 metres tall and typically inhabits primary, secondary and degraded rain forests (Versteegh and Sosef 2007). The species is widespread and common across Lower Guinea, from southern Nigeria to the western tip of Democratic Republic of Congo and is therefore ideal for studying CAR phylogeography and the nature of the climatic inversion as a phylogeographic barrier.

Here, we used a newly developed baiting kit (Couvreur et al. 2019) to sequence hundreds of nuclear markers in 112 individuals covering most of the distribution of *A. affinis*. First, we identified the major genetic clusters within *A. affinis* and their distribution to test if *A. affinis* shows a North-South genetic structuring along the climatic inversion. Second, we built a phylogenetic hypothesis of relationships among genetic clusters and conducted spatiotemporal diffusion analyses to test if lineages have frequently crossed the climatic inversion, or if dispersal has been restricted over time. We also test for an inland (eastwards) dispersal pattern, congruent with expansion out of climatically stable areas. Third, we collated information on flowering times and built ecological niche models (ENMs) of the species as a whole, as well as populations either side of the climatic inversion, to identify the underlying mechanism behind the common North-South genetic structure. Finally, we reconstruct effective population size through time to infer the past demography of each identified genetic cluster to test for recent population expansion, and if demographic histories are congruent among clusters.

## 2 MATERIAL AND METHODS

### 2.1 Sample collection

A total of 112 individuals of *Annickia affinis* were sampled across most of the species distribution range in Central Africa (Table S1). In addition, two individuals were sampled from the sister species *Annickia polycarpa* as outgroups (Couvreur et al. 2019). We obtained data on flowering and fruiting times of *Annickia affinis* using herbarium specimens of *A. affinis* retrieved from the BRAHMS database of Naturalis NL. All specimens studied in the taxonomic revision of *Annickia* (Versteegh & Sosef, 2007) were included in BRAHMS, providing a good representation of available specimens for the genus.

### 2.2 DNA extraction, gene capture and sequencing

DNA was extracted from silica gel dried leaves using the MATAB (Sigma-Aldrich, Saint Louis, Missouri, USA) and chloroform separation methods following Couveur et al. (2019). Nuclear markers were captured using the Annonaceae bait kit (Couvreur et al. 2019) made of 11,583 baits 120 bp long targeting 469 exonic regions. Barcoded Illumina libraries were constructed based on a modified protocol of Rohland and Reich (2012). See supplementary methods for details.

### 2.3 Read filtering, contig assembly and multi-sequence alignment

Reads were cleaned and filtered following the protocol in Couvreur et al. (2019) and Hybpiper (Johnson et al. 2016) was used to prepare sequence data for phylogenetic inference. Alignments were conducted using MAFFT (Katoh and Standley 2013) and cleaned with GBLOCKS (Castresana 2000). Putative paralogs for *A. affinis* that were flagged by Hybpiper were verified and removed during this process. Further information can be found in the supplementary methods.

### 2.4 SNP calling

We used SeCaPr (v1.1.4; Andermann et al. 2018) to call SNPs as it generates a psuedoreference made up of the consensus sequences for each target locus (paralogs removed) that is tailored to the given dataset, which is more efficient than the bait kit reference made from distantly related Annonaceae species. We then mapped our cleaned, paired reads to this psuedoreference using BWA (v0.7.12; Li and Durbin 2009). Duplicates were removed and SNPs were called using HaplotypeCaller in GATK (v4.0; McKenna et al. 2010). We used bcftools (v1.8; Li 2011) to apply thresholds of mapping quality (>40%) depth (>25), quality by depth (>2), minimum depth across individuals (>10) to filter SNPs. We also removed those SNPs with a minor allele frequency < 0.01, kept only biallelic SNPs and excluded monomorphic sites.

### 2.5 Population genetic clustering and statistics

We examined the genetic structure of *A. affinis* using three approaches. First, we undertook a Discriminant Analysis of Principal Components (DAPC) (Jombart et al. 2010). We used the function *find.clusters* in the R package ‘adegenet’ (Jombart 2008) to identify clusters using successive K-means with 100,000 iterations per value of k and a maximum k value of 20. We identified the most appropriate number of clusters by examining the change in Bayesian Information Criterion (BIC) with increasing values of k (number of clusters). We then used the function *dapc* in order to define the diversity between the groups identified using *find.clusters*. We performed cross-validation of our DAPC analysis to ensure our chosen number of PCs was reliable. The data were divided into training (90%) and validation (10%) sets and DAPC was carried out on the training set, with different numbers of PCs retained. 30 replicates were performed to identify the optimal number of PCs – the number that minimizes root mean squared error. We used cluster membership inferred using DAPC to define populations for downstream analyses.

Second, we used TESS3 - an approach that takes into account geographic location information when inferring population clusters (Caye et al. 2016). TESS3 was implemented using the R package ‘tess3r’ (Caye et al. 2016). We used the projected least squares algorithm and a maximum k of 20. We examined the cross-validation score for each value of K to identify the appropriate number of clusters.

Third, we used fastSTRUCTURE (Raj et al., 2014) on a reduced set of unlinked SNPs (one per locus). To maximize the number of SNPs we did not apply a depth-by-individual filter for fastSTRUCTURE analyses. We then thinned our SNPs first by applying more stringent filters (0% missing data) and then randomly choosing a single SNP from each locus, where possible. We ran fastSTRUCTURE using the default settings and the simple prior. The script ‘chooseK.py’ was used to identify the number of clusters that maximized the marginal likelihood and explained the structure in the data.

### 2.6 Phylogenetic inference

We filtered our dataset by choosing only those exons that had 75% of their length reconstructed in 75% of *A. affinis* individuals. We then used the corresponding supercontigs (i.e. targeted regions and surrounding off-target sequences) for phylogenetic inference. We added empty sequences when individuals were missing from locus alignments and we concatenated loci with the pxcat function from phyx (Brown et al. 2017). We assigned a different GTR+GAMMA model to each locus to account for differences in substitution rates. We then ran RAxML (v8.2.9) (Stamatakis 2014) using the ‘-f a’ option with 100 replicates. The tree was rooted using *A. polycarpa* as outgroup. We also conducted a coalescent-based phylogenetic analysis using ASTRAL-III (v5.5.11; Zhang et al. 2017), which uses individual gene trees to infer a species tree.

We also constructed relationships using SNPs. We constructed a phylogenetic network using splitstree (v4.14.6; Huson and Bryant 2006) and the full SNP dataset using the neighbour-net algorithm. We used the coalescent-based Bayesian phylogenetic approach SNAPP (Bryant et al. 2012) in BEAST2 (v2.5.2; Bouckaert et al. 2019) to infer relationships among different genetic clusters. We used the same SNP dataset as used with fastSTRUCTURE. We randomly chose 10 individuals per DAPC genetic cluster (or all individuals if a cluster had <10) as representatives. We ran the chain for 1 million generations, sampling every 1000 generations until effective sample size (ESS) values, assessed with Tracer v1.7 (Rambaut et al. 2018), were over 200 for all parameters. We performed two runs and combined these to ensure that they reached stationarity at the same point. We then generated a consensus tree using TREEANNOTATOR (Bouckaert et al. 2019).

### 2.7 Phylogeographic Diffusion in Continuous Space

We reconstructed the spatiotemporal dynamics of *A. affinis* using BEAST v1.8.4 (Drummond and Rambaut 2007) and spreaD3 v0.9.6 (Bielejec et al. 2011). As this analysis is computationally intensive we used a subset of our dataset by selecting only the five most informative loci based on number of phylogenetic informative sites. We added a partition consisting of longitude and latitude coordinates as continuous trait data. We used a HKY+G substitution model and a strict clock model for each genetic locus and an exponential growth coalescent tree prior. We ran the analysis for 100 million generations and assessed ESS values using Tracer v1.7 (Rambaut et al. 2018). We then used spreaD3 to visualize the output at several time points during the history of *A. affinis*. We repeated this analysis with the next five most informative loci to ensure similar patterns were recovered across datasets.

### 2.8 Demographic history

We used stairway plot (v2; Liu and Fu 2015), a model-flexible approach that uses site frequency spectra (SFS) to infer changes in effective population size (*N*_*e*_) through time. We generated filtered VCF files representing each cluster as detailed above but did not apply a minor allele frequency filter. We then calculated folded SFS for each cluster. Stairway plot uses SNP counts to estimate the timing of events and changes in *N*_*e*_ so the removal of SNPs with missing data may skew counts. To overcome this we used an approach documented in Burgarella et al. (2018) where we modified each SFS by first calculating the minor allele frequency at each SNP and then multiplying this by the mean number of sequences (haploid samples) at each site. This results in a new SFS that makes use of all observed site frequencies and minimizes the number of SNPs removed. The total of samples is slightly reduced based on the amount of missing data. The number of random breakpoints were calculated as recommended in the manual. We used 67% of sites for training and performed 200 bootstrap replicates. The number of observed sites was calculated as the total length of the pseudoreference. We used an angiosperm wide mutation rate of 5.35 × 10 ^−9^ sites/year (De la Torre et al. 2017) and a generation time of 15 years based on the generation time of the Annonaceae species *Annona crassiflora* (Collevatti et al. 2014). In addition, sequencing error can skew the SFS by inflating the number of singletons. We ran analyses using the entire SFS, and then reran with singletons removed to ensure similar histories were reached. We also ran analyses with a generation time of 50 years (Baker et al. 2014) to examine the effect this increase had on population size change.

### 2.9 Modelling of current and past ranges

Current and Last Glacial Maximum (21k years ago; LGM) potential distributions were modelled using MaxEnt (v3.3.3; Phillips et al. 2006) as implemented in ‘biomod2’ (v3.3-7.1;Thuiller et al.2009). Current and LGM (MIROC global circulation model) climatic data were downloaded from WordClim ver. 1.4 (Hijmans et al. 2005) at a resolution of 10*10 arc-minute. The LGM period represents the latest unfavourable climate for tropical species and is therefore a good period to model the impact of past climate change on potential range. A total of 346 presence data points (Table S1) covering the known distribution of *A. affinis* were spatially filtered to one point per cell to avoid overfitting due to sampling bias. Model overfitting was constrained by using the ⍰ regularization parameter in Maxent, which limits model complexity (Radosavljevic and Anderson 2014), and was set to 2.00 and 4.00, rather than the default MaxEnt value of 1.00. Modelling with all 19 bioclim variables produced unrealistic results and failed to properly model the current species range independent of the regularization parameter (results not shown). Using just eight bioclim variables (four precipitation and four temperature, see supplementary methods) greatly improved the accuracy of the models to the known distribution. Model performance was evaluated using a cross-validation procedure (Ponder et al. 2001, Muscarella et al. 2014, see supplementary methods). Model fit was assessed using area under curve (AUC; Elith et al. 2006) and the true skill statistics (TSS, Allouche et al. 2006). The best fitting model was then projected into the LGM.

We then constructed ecological niche models (ENMs) splitting them into two groups, either North or South of the climatic inversion (above or below 2°N). We compared these models using the R package ‘ENMTools’ (Warren et al. 2019). Models were built using the generalized linear model (GLM) approach using the same bioclimatic variables that were used in our MaxEnt models above. We performed niche equivalency tests (100 replicates) using the metrics Schoener’s *D* (Schoener 1968), *I* (Warren et al. 2008) to assess whether ecological niches of individuals North and South of the climatic inversion were more different than expected by chance. We also performed symmetric background tests (100 replicates; Warren et al. 2008), to correct for habitat availability. If observed values of *D* or *I* are significantly different than the simulated distribution we can state whether the observed similarity between populations is significantly larger or smaller than expected in the context of the available environment. We then assessed ecological niche overlap in a phylogenetic context using our SNAPP tree (100 replicates; Warren et al. 2008).

## 3 RESULTS

### 3.1 Sequencing

A total of 124.7 million reads were generated for 112 *A. affinis* individuals at an average coverage depth of 77.5x across all targeted loci. Using HybPiper we identified 366 loci where 75% of the exon length was recovered in at least 75% of individuals. A total of 15 loci showed signs of paralogy and were removed, leading to a final dataset of 351 supercontigs totalling 756 kb of sequence data. After cleaning and filtering our SNP calling approach yielded 5,964 high-quality SNPs from 262 different loci. A comparison of the output of the SeCaPr and HybPipier pipeline include amount of genetic variation and length of sequence recovered can be found in Table S2. We found 240 loci that were reconstructed in both pipelines, representing 78% of the total number of SeCaPr loci (306) and 68% of HybPiper loci (351).

### 3.2 How are populations structured across the range of *A. affinis*?

After cross-validation the number of axes that was associated with the lowest associated root mean squared error was 80 so this value was used in the DAPC. Changes in BIC greatly decreased after k = 4 (Fig. S1) suggesting that four clusters best fit our data (Fig. 1, Fig. S2). Two major clusters contained 35 and 63 individuals that were located primarily in Western Gabon (cluster WG) and Cameroon (cluster CA) respectively. Two smaller clusters of seven individuals each were located in eastern Gabon (cluster EG) and Gabon / Republic of Congo (cluster GC). There is a clear discontinuity in genetic structure across the equator, separating cluster CA from the rest, except for a pair of individuals belonging to cluster EG.

The TESS3 analysis also found that four clusters best defined our data (Fig. 1B, S3) with geographic discontinuities generally congruent between analyses (Fig. 1; Fig. S4). Individual admixture proportions revealed generally limited mixed ancestry within samples (Fig. 1B), though we detected a conspicuous amount at a single location. The two northern most individuals belonging to cluster EG, found in at Meyo Centre in Cameroon (labelled in Fig. 1A), had a considerable proportion of their ancestry from cluster CA (Fig. 1B). The inverse was true of the two individuals from cluster CA that were from the same location. The fastSTRUCTURE analysis was similar to the aforementioned analyses, even with a reduced SNP dataset, though the grouping of individuals belonging to DAPC clusters GC and EG was not consistent for different values of K (Fig. S5). However, there was little evidence for admixture when using this approach including for “Meyo Centre” individuals. These differences are likely because of the relatively small numbers of SNPs used. Our inferred phylogenetic network (Fig. 1C) also revealed four major clusters, and that clusters WG & CA and EG & GC were grouped together.

### 3.3 How did populations of *A. affinis* disperse across central Africa?

The ASTRAL tree (Fig. 1D) was generally well-supported at deeper nodes with high values of local posterior probability. This suggests that there are a high proportion of gene trees supporting the topology in figure 1D, making it a reliable estimation of evolutionary history among major clades. The tree topology reflected our clustering inferences, lending further support to our four identified clusters and robust evidence for phylogeographic structuring. Our SNAPP analyses reach stationarity after less than 1 million generations and separate runs were combined as they converged at the same point. The same pattern of divergence inferred for the four genetic clusters using ASTRAL was found in our RAxML tree (Fig. S6) and our SNAPP tree (Fig. S7). The phylogenetic network (Fig. 1C) also yielded relationships that supported two groups of clusters: CA & WG and EG & WC.

We assessed the geographic locations in these clusters at a finer scale by mapping each tip of the ASTRAL phylogenetic tree to its collection site (Fig. S8). In cluster WG (Fig. S8C) the first of two major clades contains individuals that are found near Mt. Cristal and in coastal rain forests in Northwest Gabon. The remaining individuals in cluster WG formed a monophyletic group and are found to the South and East, as far as the Southern tip of the Republic of Congo. We then examined the geographic locations in cluster CA and identified three clades with distinct geographic distributions (Fig. S8D) going up Cameroon’s Atlantic coast. Individuals belonging to the largest clade extend inland. Given this structure we repeated DAPC clustering analyses using only individuals from cluster CA and revealed fine-scale genetic structure that supported these three clades (Fig. S9).

Our diffusion analysis was based on 47.3kb of sequence data across five partitions and converged with the ESS > 200 for all parameters after 100 million generations. The root was inferred to be around central Gabon (Fig. 2A). We estimated a single lineage crossed the climatic inversion, from South to North, establishing cluster CA (Fig. 2). Late in the evolutionary history of *A. affinis* another dispersal event crossed the barrier at Meyo Centre (see Fig. 1A). Our repetition of the diffusion analysis with different loci matched these patterns (Fig. S10) indicating our results are reliable and unlikely to have been biased by particular gene histories.

**Fig. 2.**
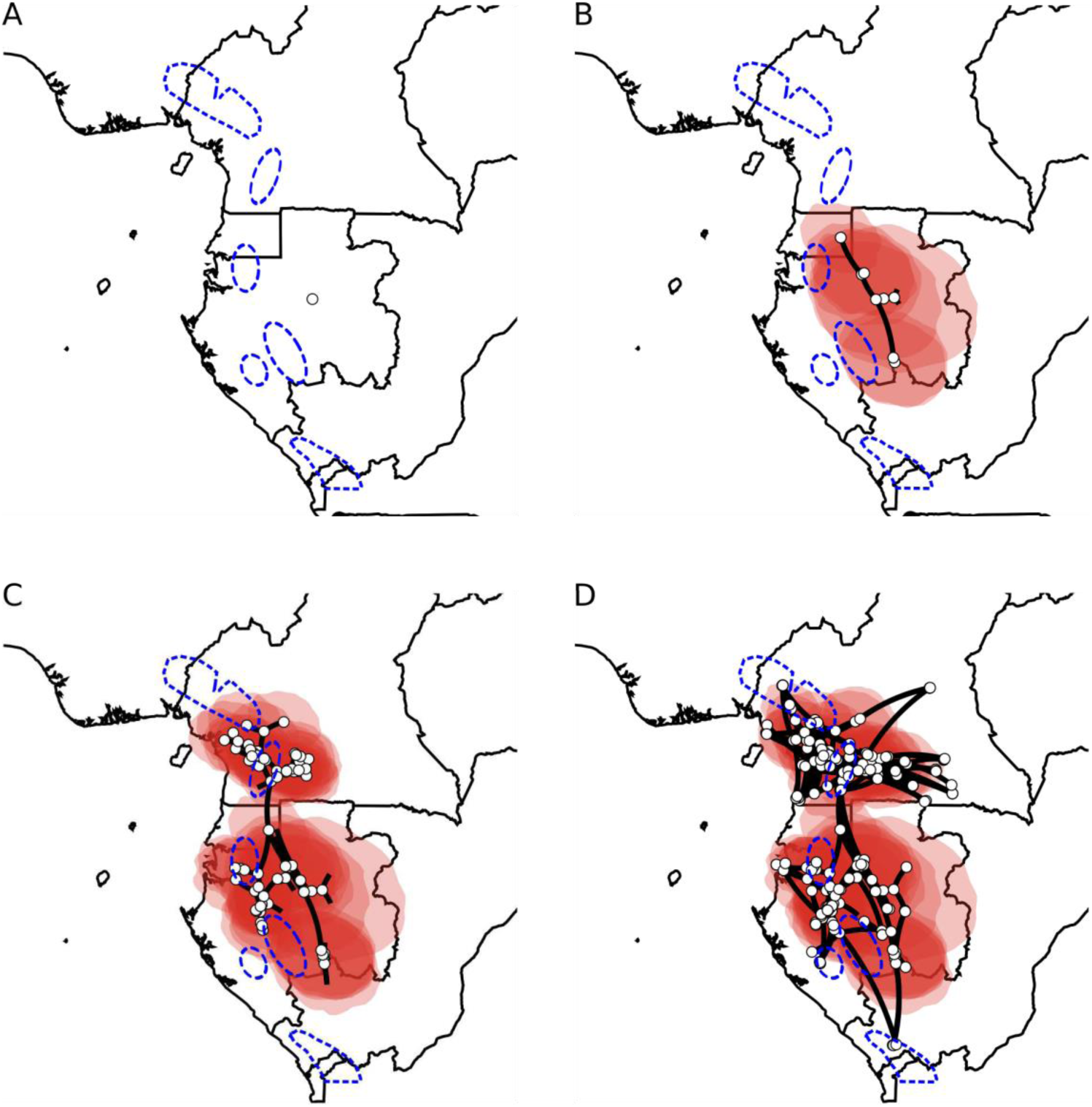
Phylogeographic diffusion analysis split into four time slices. Images were rendered using spreaD3 and move forward through time starting from the (uncalibrated) time of the most recent common ancestor (A) to the present day (D). White circles represent ancestrally estimated geographic locations for nodes in the inferred phylogenetic tree, as well as current, real locations at tips. Polygons around points represent uncertainty of estimated ancestral locations at 80% highest posterior density (HPD). Putative refugia following Maley (1996) are shown in dashed blue lines.

### 3.4 Do different populations share similar demographic histories?

We estimated the demographic history of the four DAPC clusters (Fig. 3; Fig S11-14 for full plots). Over the last 100 thousand years (Ka) three clusters (GC, WG and CA) experienced similar demographic histories with population decline around 50-70 Ka followed by an increase in *N*_*e*_ towards the present. We found this increase began at slightly different times across these three clusters though all show rapid increases in population size close to the LGM, around 20-27 Ka. Cluster CA showed evidence of a rapid growth very close to the present, in the last 5 Ka. Cluster EG had a very different history, exhibiting a relatively constant population size in the past with a gradual decline in the last 10 Ka. Results were very similar when singletons were removed indicating that sequencing errors were not affecting our analyses (results not shown) and increasing the generation time to 50 years had little effect on the timing and trend in population size change but did noticeably decrease *N*_*e*_ (Fig. S15).

**Fig. 3.**
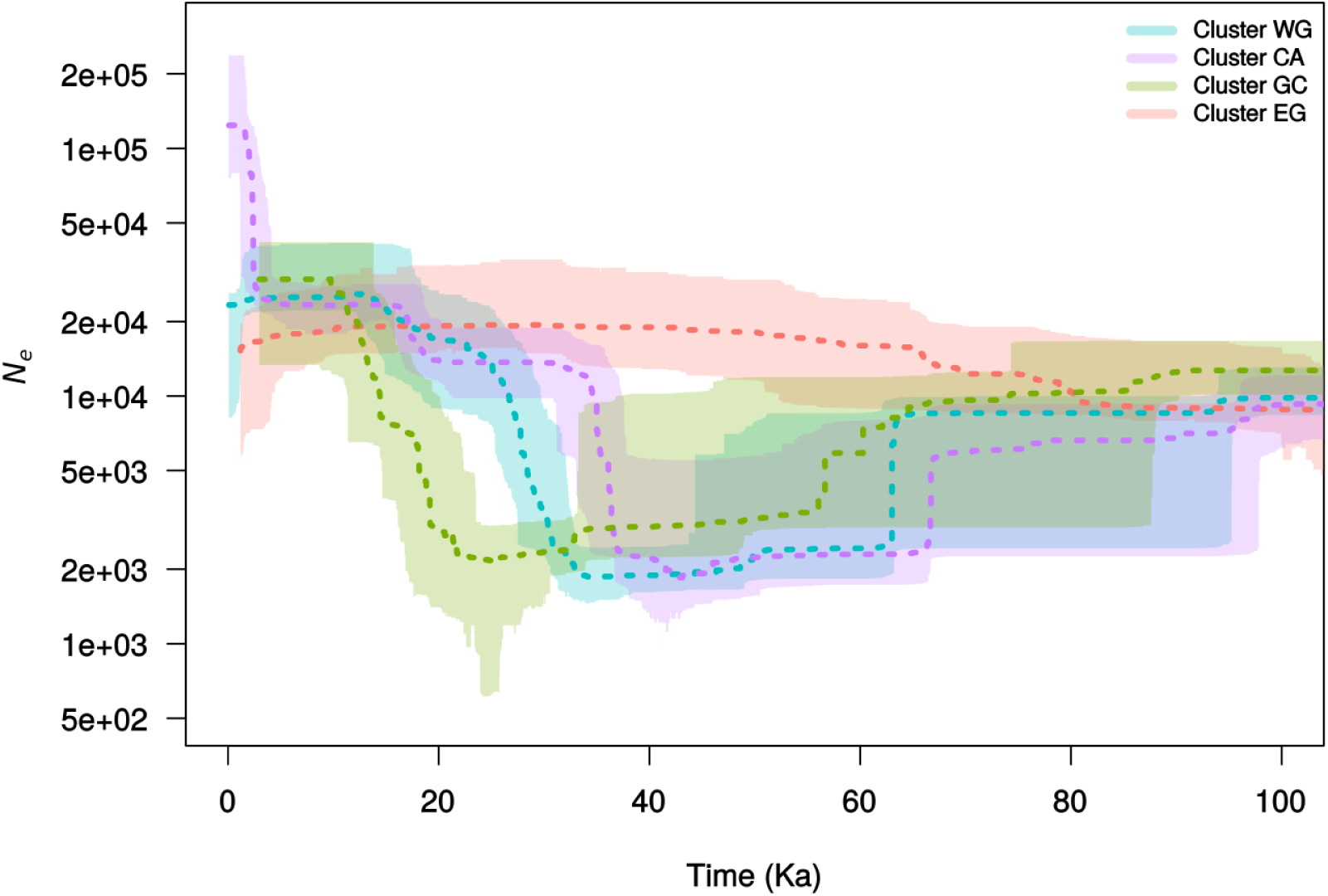
Plots of effective population size through time for each of the four clusters inferred using stairway plot. The present is located on the left side of each graph. The dotted line represents the median population size and the shaded polygon represents the 80% central posterior density intervals. Colours correspond to the colours used in figure 1. Full plots of each species can be found in figures S11-14.

### 3.5 Which areas have remained climatically stable over time?

A total of 113 data points were retained after filtration. The best predictors were Precipitation of Wettest Month (Bio13) and Precipitation Seasonality (Bio15) (Table S3). A regularization multiplier of 2 generated a better model fit than with 4, showing a good visual match with the known distribution of the species at present (Fig. 4A). The mean value of the AUC for the training and test data were respectively 0.77 and 0.76. The mean value of TSS was 0.454, indicating that the model is better than a random model. During the LGM, the highest presence probabilities were all located along the Atlantic coast in Cameroon, Equatorial Guinea and Gabon (Fig. 4B).

**Fig. 4.**
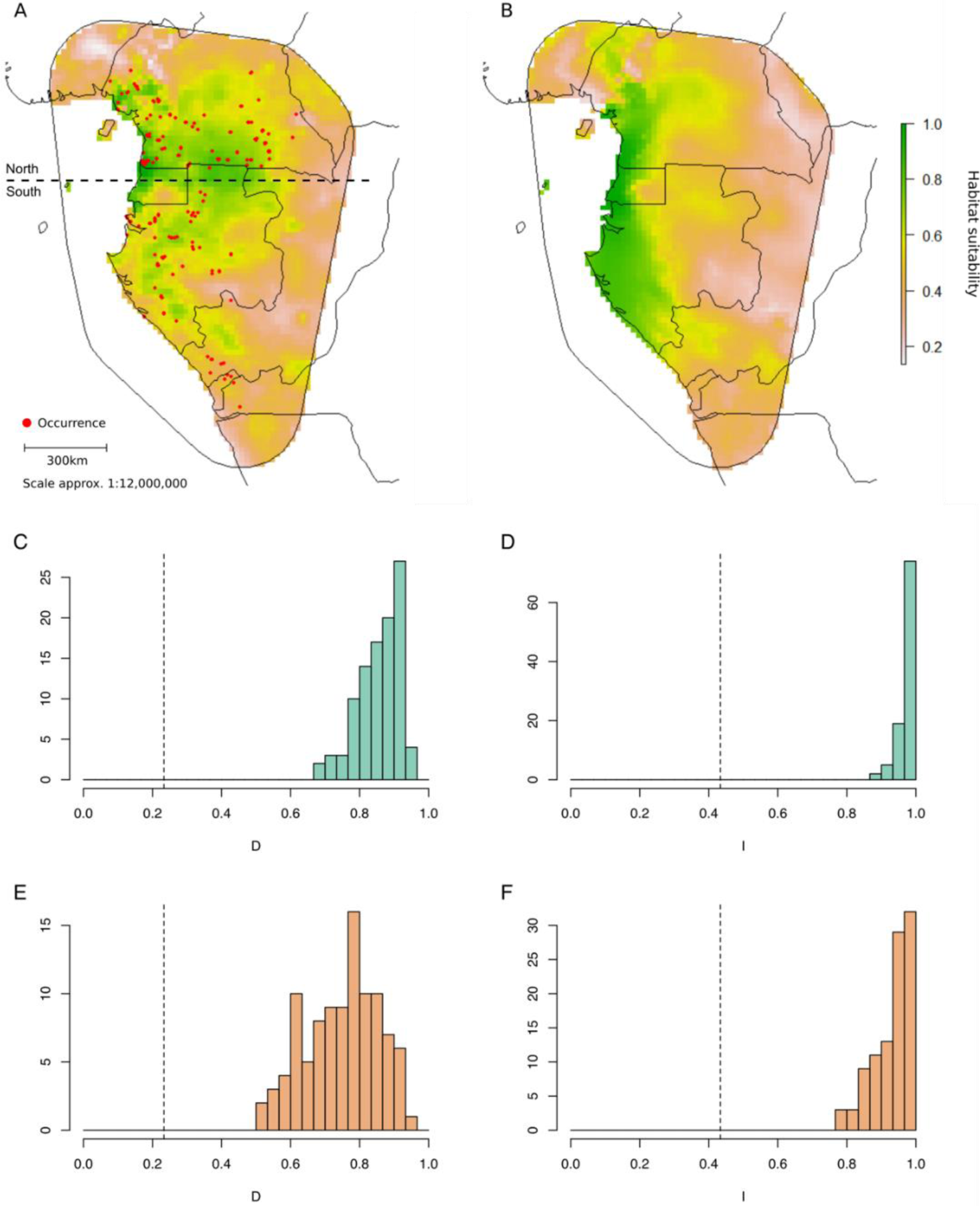
Ecological niche models (ENMs) for the present (A) and projected into the past, during the last glacial maximum (B). ENMs were constructed using MaxEnt and bioclimatic variables. The colour scale represents habitat suitability for each cell where green indicates more suitable cells. Red circles in (A) indicate sites where *A. affinis* individuals were collected and used in building the model. Comparisons of ENMs constructed from individuals North and South of the climatic inversion (see panel A) are shown in panels (C-F). Panels (C) and (E) show histograms of 100 simulations of identity tests using two different metrics, *D* and *I*. Similarly, (E) and (F) are background tests, taking into account the available environments when comparing models. Values of *D* and *I* range from 0 (no niche overlap) to 1 (identical niches among the two groups). Dotted lines indicate observed values.

### 3.6 Does flowering time or ecological niche diverge across the climatic inversion?

We collated fruiting and flowering time data from 18 individuals (Table S4), 15 of which had dates associated, and plotted their distribution by month (Fig. S16). This revealed little change in flowering or fruiting time either side of the inversion. Flowering and fruiting appears to occur all year round in the North while in the South the majority of these events occurred from September to January (though individuals here were also found to flower in April).

Comparisons of ENMs for individuals North and South of the climatic inversion revealed large differences in ecological niches among these geographically separated groups. Niche identity tests indicated that there are significant differences in niche space of individuals North and South of the inversion (p < 0.01; Fig. 4C, D). Our symmetrical background tests (Fig. 4E, F) revealed that correlation between the available habitat between these areas is moderate-high, but niche overlap is significantly lower than would be expected based on available habitat (p < 0.05). When we examined niche overlap in a phylogenetic context, we found that overlap was smallest between clusters CA and WG (Fig. S7A), which corresponds to divergence across the climatic inversion.

## 4 DISCUSSION

### 4.1 Limited dispersal and divergent niches across the climatic hinge

Intraspecific diversity, based on phylogenomic nuclear sequence data, within the widespread tree species *Annickia affinis* is highly structured with a clear North-South divide between identified genetic clusters (Fig. 1). This is the first time this has been observed in plants using genomic data and adds to the growing evidence of an important phylogeographic barrier around a climatic hinge between 0-3°N in numerous CAR distributed plants (Hardy et al. 2013; Heuertz et al. 2014; Faye et al. 2016; Ley et al. 2017; Pineiro et al. 2017). This North-South discontinuity is, however, generally not recovered in CAR distributed animals except in rare cases (e.g. Portik et al. 2017). This suggests that the processes taking place in relation to this barrier affect the flora of CARs more than the fauna. Indeed, Blatrix et al. (2017) showed that this barrier was more abrupt in the studied tree species (*Barteria fistulosa*) than within the associated symbiotic ants. However, the reasons for this genetic break in a seemingly continuous rain forest region remain little understood (Hardy et al. 2013).

Here, we show that, throughout the evolutionary history of *A. affinis*, a single major northward cross-hinge colonisation event occurred leading to the successful establishment of the Cameroonian population (Fig. 2C). Hardy et al. (2013) posited that environmental differences either side of the inversion may limit or prevent successful dispersal and establishment. The rarity of lineage dispersal across the hinge that we observed lends support to this hypothesis. It may not be environmental constraints that limit dispersal, but instead the mechanism of dispersal itself. However, the small red to black fleshy fruits of *Annickia affinis* (Versteegh & Sosef, 2007) are frugivore-dispersed (Poulsen et al. 2001; Holbrook and Smith 2000) and can potentially travel long distances (> 500 m) for example by hornbills. Thus, the genetic structure of *A. affinis* in general, and the North-South divide in particular, is not linked to seed dispersal limitation *per se*. Our ENM comparisons give further support to the environmental differences hypothesis - we found that ecological niches were significantly different between individuals North and South of the inversion (Fig. 4C-F). Furthermore, when viewed in a phylogenetic context, ecological niches of clusters CA and WG showed much smaller values of niche overlap than values at other nodes in our population-level tree (Fig. S7A).

We detected that one genetic cluster (cluster EG) extends across the climatic hinge into south Cameroon (Meyo Centre site, Fig. 1A) leading to a second, more recent, northwards migration event into the climatic hinge area (Fig. 2D). This indicates that the barrier is not absolute, agreeing with other studies (Hardy et al. 2013; Duminil et al. 2015; Pineiro et al. 2017). The Meyo Centre site lies within the inversion zone and several individuals with mixed ancestry are found here, including those in the second dispersal event (Fig. 1A, B). A similar result was found in *B. fistulosa*, with 20% of individuals sampled near 1⍰N being backcrossed or F_1_ individuals (Blatrix et al. 2017). Aside from these few individuals, we found a general lack of evidence for admixture between genetic clusters on either side of the inversion (Fig. 1B; Fig. S5), indicating that gene flow is generally rare.

The hypotheses of Hardy et al. (2013) were not mutually exclusive and given that there is no clear barrier to dispersal of pollen or seeds, divergence in traits such as flowering time could play a role in preventing reproduction and limiting North-South admixture. This may in turn be linked to why successful establishment across the inversion is rare. However, we observed no clear difference in flowering time among individuals north and south of the climatic inversion (Fig. S16). We note that data was available for just15 individuals, so more is needed to reliably assess whether flowering time differs.

Interestingly, similar north-south phylogeographic breaks are also known from the Atlantic rain forests of Brazil, due to differing climatic regimes and floral compositions (Carnaval et al. 2014; Leite et al. 2016). This suggests that similar processes, though not necessarily driven by exactly the same factors, might be driving patterns of intraspecific diversity in different TRF regions.

### 4.2 Out-of-refugia migration in northern forests

The refuge hypothesis has received support from several population genetic studies of CAR plants, showing concordance between putative refugia and regions of high or unique allele/haplotype diversity (Lowe et al. 2010; Dauby et al. 2014; Heuertz et al. 2014; Faye et al. 2016). Here we find that the inferred evolutionary dynamics of *A. affinis* support a potential role for Pleistocene forest oscillations in shaping intraspecific genetic diversity patterns, but this role may not have been the same in each region. we found evidence for recent demographic expansion in three clusters (CA, GC and WG, Fig. 3), as would be expected if *A. affinis* expanded out of refugia. These expansions were estimated to have over the last 40 Ka, around the timing of the LGM. This timing was fairly robust to changes in generation time used (Fig. S15). However, further work is needed to determine a more accurate mutation rate and generation time for *A. affinis* to verify the timing of these events, and we would therefore caution against over-interpreting the exact timing of demographic events and instead focus on the population size trends. Sampling sizes were also small (n =7) for clusters GC and EG meaning we are less confident in the patterns reconstructed for these clusters. Similar patterns of recent expansion were detected in populations of central African plants (Pineiro et al. 2017) and animals (Bell et al. 2017) as well as in studies on neotropical flora (Vitorino et al. 2016) and fauna (Batalha-Filho et al. 2012).

In contrast to the other clusters, EG showed constant population size with a slight decline towards the present, perhaps indicating that glacial cycles have not played an important role in its demographic history. Similar demographic patterns were found in populations of two central African *Erythrophleum* species (Duminil et al. 2015), though these exhibited a more pronounced decline in the last 50 Ka. Overall, our results indicate that demographic responses to past climate change have been different among populations of *A. affinis* across central Africa. Similar patterns of recent expansion were detected in populations of central African plants (Pineiro et al. 2017) and animals (Bell et al. 2017) as well as in studies on neotropical flora (Vitorino et al. 2016) and fauna (Batalha-Filho et al. 2012).

We also find evidence to support the scenario presented by Anhuf et al. (2006) who proposed that coastal rain forests in central Africa acted as refugia during the LGM, though results again vary among regions. The modelled LGM distribution of *A. affinis* indicates that suitable habitat was concentrated continuously along the coast, from Cameroon to Gabon (Fig. 4B), like in the understory palm species *Podococcus barteri* (Faye et al. 2016). In addition, we uncovered fine-scale genetic structure and evidence for within-population dispersal eastwards in Cameroon (Fig. 2), demonstrating a possible out-of-refugia pattern in this area. By contrast, we did not find an inland pattern of migration in Gabon where dispersal was both eastwards and westwards from a more central area. This may be a result of there having been a large area with highly-suitable conditions (>0.8) during the LGM that extended further from the coast in Gabon than in Cameroon (Fig. 4), meaning that populations could persist and expand out of this area. In addition, we inferred more pronounced East-West clustering (Fig. 1) in Gabon than in Cameroon, which has been observed in at least four other CAR tree species (Hardy et al. 2013).

The spatial diffusion approach has been shown to recover the true ancestral locations under a generative model that is the same as used for inference (Lemey et al. 2010). Related categorical approaches known as “mugration” discrete trait analysis (DTA) have received criticism and been largely replaced by structured coalescent approaches (De Maio et al. 2015). These issues may be applicable to continuous diffusion but no equivalent spatially-explicit coalescent implementation exists. As for any model, violations of its assumptions need to be considered and sampling bias is likely to be an issue for reconstructing the history of migration (Lemey et al. 2014; De Maio et al. 2015). The ways that this may affect a continuous mode are not well characterised and warrant further work. Given that our populations, in particular EG and GC, may have been underrepresented we must interpret our results with caution. Finally, the four retrieved clusters are found in allopatry or parapatry (Fig. 1A). This supports the hypothesis of incomplete mixing after post-glacial expansion and is similar to patterns found in other CAR species (Hardy et al. 2013) and within species from other TRF regions (Carnaval et al. 2009; Leite and Rogers 2013). Bringing our results together, it appears that refugia may have played a different role for populations in different areas, and that each has responded to past climate in change in its own way.

## 5 CONCLUSIONS

This study uncovered the evolutionary dynamics and demographic history of the CAR tree species *Annickia affinis*. Our approach is the first to use genome-wide data from hundreds of nuclear loci to infer population-level phylogeographic patterns in CAR plants. We highlighted how a climatic inversion limits lineage dispersal, drives ecological niche divergence and shapes patterns of population structure across a continuous rain forest region. We also show that the current distribution of extant populations is the result of different demographic histories and, in northern regions, migration eastwards from putative refugia in coastal rain forests. Our study is an example of how taking advantage of recently developed genomic resources, such as the sequence capture kit used here, can help to improve our understanding of TRF evolution, at the population level and above.

## ACKNOWLEDGEMENTS

We thank Prof. Moutsambote, Raoul Niangadouma, Théophile Ayole for help in the field. The authors acknowledge the IRD itrop HPC (South Green Platform) at IRD Montpellier for providing HPC resources. We are grateful to the Centre National de la Recherche Scientifique et Technique (CENAREST), the Agence National des Parques Nationaux (ANPN) and Prof. Bourobou Bourobou for research permits (AR0020/16; AR0036/15 (CENAREST) and AE16014 (ANPN). Fieldwork in Cameroon was undertaken under the “accord cadre de cooperation” between the IRD and Ministère de la Recherche Scientifique et Technique (MINRESI). Prof. Bouka Biona of the Institut National de Recherche en Sciences Excates et Naturelles (IRSEN) of the Republic of Congo is thanked for research permits. Version 3 of this preprint has been peer-reviewed and recommended by Peer Community In Evolutionary Biology (https://doi.org/10.24072/pci.evolbiol.100094).

## Data availability

The fastq (R1 and R2) sequences for all individuals are available in Genbank SRA under Bioproject number PRJNA508895 (http://www.ncbi.nlm.nih.gov/bioproject/508895). Scripts for bioinformatics can be found at https://github.com/ajhelmstetter/afrodyn.

