## supplementary materials for "Phylogenomic approaches reveal how a climatic inversion and glacial refugia shape patterns of diversity in an African rain forest tree species"

ELECTRONIC SUPPLEMENTARY MATERIAL

TABLE OF CONTENTS

**Appendix S1** Supplementary materials and methods

**Figure S1** BIC values for different numbers of clusters

**Figure S2** DAPC scatterplot

**Figure S3** Cross-validation for TESS3 analyses

**Figure S4** Map of TESS3 genetic clusters

**Figure S5** fastSTRUCTURE barplots

**Figure S6** ASTRAL and RAxML phylogenetic trees

**Figure S7** SNAPP analysis and phylogenetic ENM overlap

**Figure S8** ASTRAL tree split by cluster and mapped to sample locations

**Figure S9** DAPC for cluster CA only

**Figure S10** Repetition of diffusion analysis

**Figures S11-S14** Stairway plots for genetic clusters with confidence intervals

**Figure S15** Stairway plots using a generation time of 50 years

**Figure S16** Flowering and fruiting times

**Table S1** Specimen data table

**Table S2** Comparison of HybPiper and SeCaPr pipelines

**Table S3** Maxent variable contributions

**Table S4** Phenological data

**Appendix S1 SUPPLEMENTARY MATERIALS AND METHODS**

**1.1 Library preparation and sequencing**

Libraries were prepared using 6-bp barcodes and Illumina indexes to allow for multiplexing at different levels. Briefly, total DNA for each individual was sheared using a Bioruptor Pico (Diagenode, Liége, Belgium) to a mean target size of 500 bp. DNA was then repaired, ligated and nick filled-in before an 8–11 cycle prehybridization PCR was performed. After clean-up and quantification, libraries were bulked, mixed with biotin-labeled baits and hybridized to the targeted regions using the bait kit designed above. The hybridized biotin-labeled baits were then immobilized using streptavidin-coated magnetic beads. A magnetic field was applied and supernatant containing unbounded DNA was discarded. Enriched DNA fragments were then eluted from the beads and amplified in a 14–16 cycle real-time PCR to complete adapters and generate final libraries. Libraries were sequenced on an Illumina HiSeq v3 platform pair end and length of 150 bp (Illumina, SAn Diego CA, USA) at CIRAD facilities (Montpellier, France) with around 18 pmol of the capture-amplified DNA libraries deposited on the flowcell.

**1.2 Bioinformatics**

We demultiplexed with a 0-mistmatch threshold using the demultadapt script (<https://github.com/Maillol/demultadapt>) and adapters were removed using cutadapt 1.2.1 (Martin, 2011) with the default parameters. Read quality was filtered according to their length (>35pbp) and quality mean values (Q > 30) using a custom script (https://github.com/SouthGreenPlatform/arcad-hts/blob/master/ scripts/arcad_hts_2_Filter_Fastq_On_Mean_Quality.pl). Forward and reverse sequences were paired according to their name in the fastq files using a comparison script, adapted from TOGGLe (Tranchant-Dubreuil et al., 2018). A terminal trimming of 6 bp was performed on reverse sequences to ensure removal of barcodes in case of sequences shorter than 150 bp using the fastx trimmer script which is part of the fastx toolkit (<https://github.com/agordon/fastx_toolkit>).

**1.3 Contig assembly and sequencing alignment**

Briefly, HybPiper (v1.2) (Johnson et al., 2016) was used to process our data, identifying target exonic regions as well as off-target introns. This pipeline yields ‘supercontigs’ containing target and off-target sequence data. Intronic sequences are typically more variable than exon sequences and it is therefore useful to incorporate when inferring relationships between recently diverged taxa. We aligned supercontigs corresponding to recovered target exons using MAFFT (v7.305) (Katoh and Standley, 2013) with the "--auto" option and cleaned these alignments with GBLOCKS (v0.91b) (Castresana, 2000) using the default parameters and all allowed gap positions.

HybPiper flags potentially paralogous loci and we assessed those loci by building gene trees including all putative paralogs. If these gene tree grouped by paralog (i.e. potential paralogs are more closely related to paralogs in other taxa than the alternative sequences in the same taxon) we designated the locus as a true paralog and removed it from downstream phylogenetic inference. Paralogs were identified using only *A. affinis* individuals.

**1.4 Genetic clustering**

We visualised fastSTRUCTURE results using the R package ‘pophelper’ (Francis, 2016). Various other R packages ‘adegenet’ (Jombart 2008), ‘pegas’ (Paradis 2010), ‘poppr’ (Kamvar et al. 2014) and ‘hierfstat’ (Goudet 2005) were used in the course of our population genetic analyses.

**1.5 Species distribution modelling**

Location data were filtered spatially to one point per cell to avoid overfitting due to sampling bias. This resulted in a total of 113 grid cells of 10 * 10 arc-minutes. All climate data were downloaded from WordClim ver. 1.4 (Hijmans et al. 2005) at a resolution of 10 * 10 arc-minute. Eight bioclim variables were selected for this analysis: Annual Mean Temperature (Bio1), Mean Temperature of Warmest Quarter (Bio10), Mean Temperature of Coldest Quarter (Bio11), Annual Precipitation (Bio12), Precipitation of Wettest Month (Bio13), Precipitation of Driest Month (Bio14), Precipitation Seasonality (Bio15), Precipitation of Wettest Quarter (Bio16). Past climate data for the LGM (21,000 years ago) was estimated by climatic projections of the MIROC global circulation model (GMC) following Faye et al. (2016). Indeed, the MIROC is the only GCM predicting a reduction of precipitation across central Africa during the LGM as expected. We evaluated model performance using a cross-validation procedure. Instead of absence data, pseudo-absence data were generated using the RAINBIO database (Dauby et al. 2016) following the “Target group sampling” method (Ponder et al. 2001). Presence and pseudo-absence data were split into training and test datasets. We used the checkerboard method (Muscarella et al. 2014) to separately test and train data so that spatial structure of data was maintained. Several test and training data sets were created, and we assessed each independently. The performance of the models was estimated by calculating the area under curve (AUC), a threshold-independent measure of performance (Elith et al. 2006) and the true skill statistic (TSS, Allouche et al. 2006) for each pair of presence and pseudo-absence data. TSS values range from -1 to +1, where +1 is an indication of a perfect model fit and values ≤ 0 is an indication that models which are no better than random (Allouche et al., 2006).

SUPPLEMENTARY FIGURES

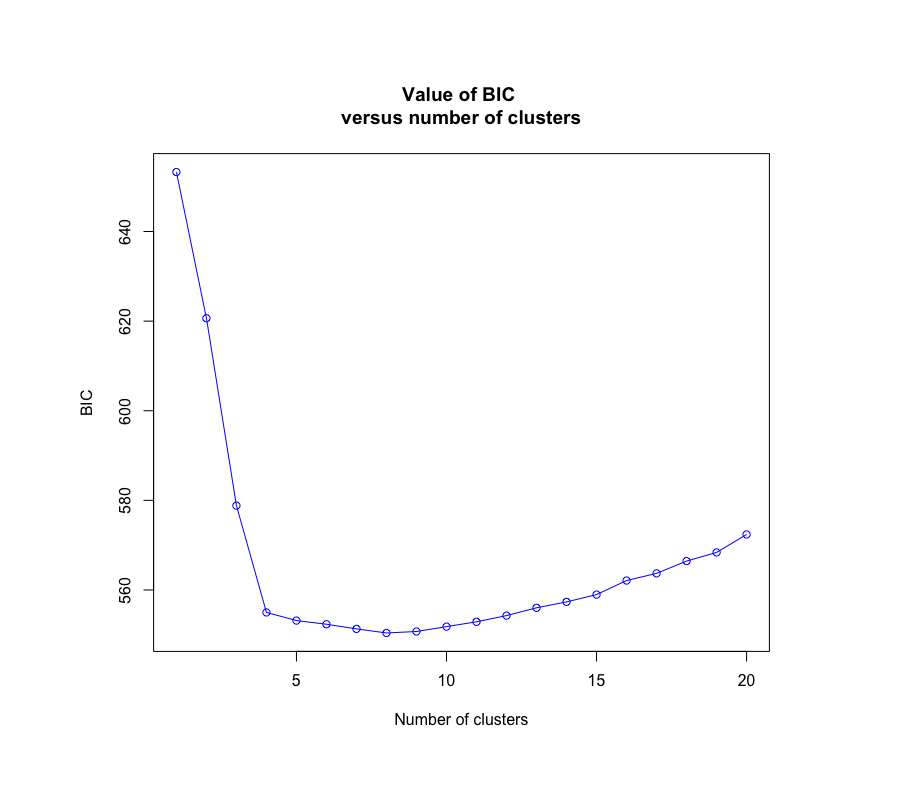

**Fig. S1** Number of clusters (k) was chosen by examining how BIC changed with increasing values of k. Four clusters were chosen because of the rapid drop in the change in BIC between four and five clusters.

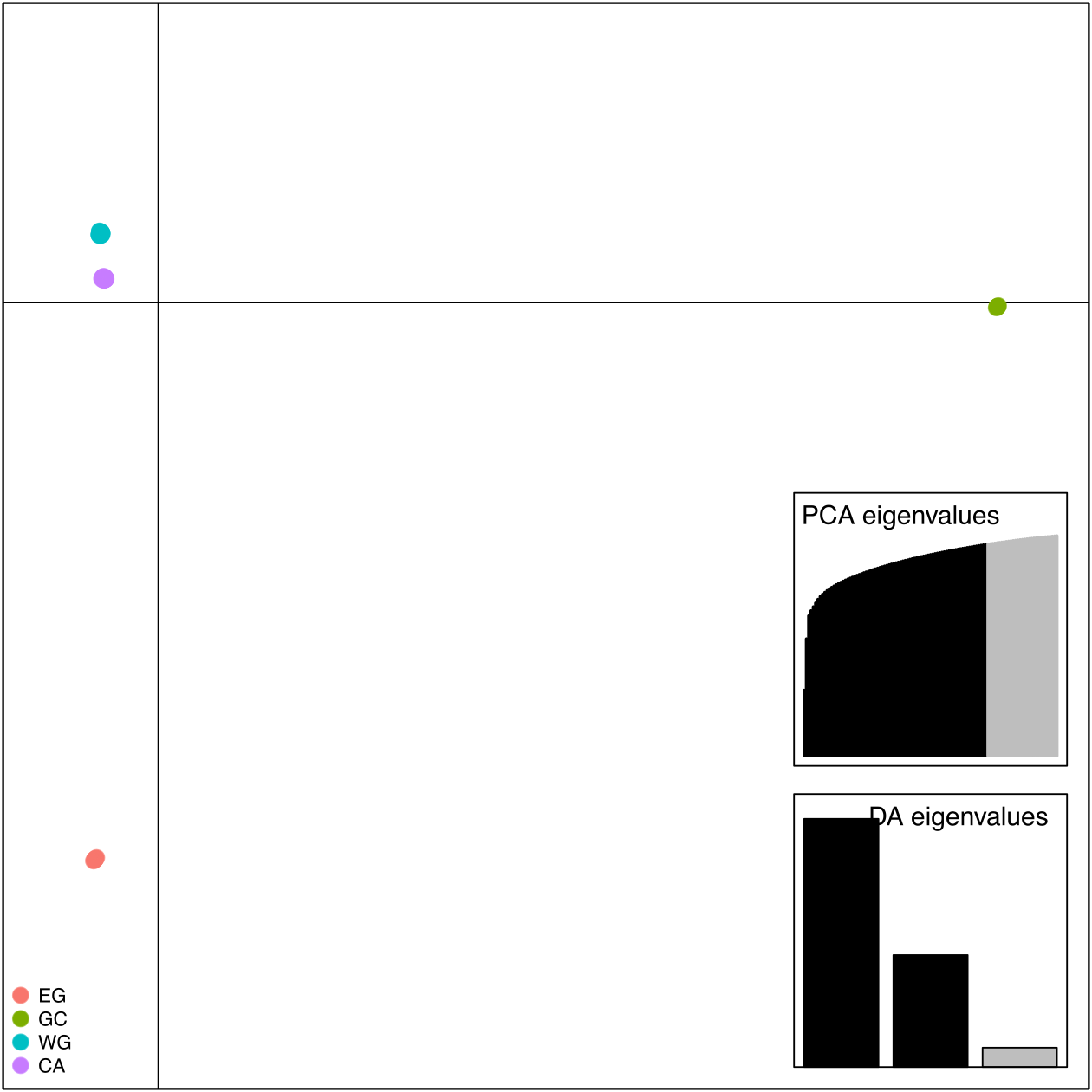

**Fig. S2** Scatterplot of DAPC analyses. Points represent individuals and inferred clusters (k = 4) are coloured as in Fig. 1. Inset are screen plots of PCA and DA eigenvalues where black shows retained eigenvalues.

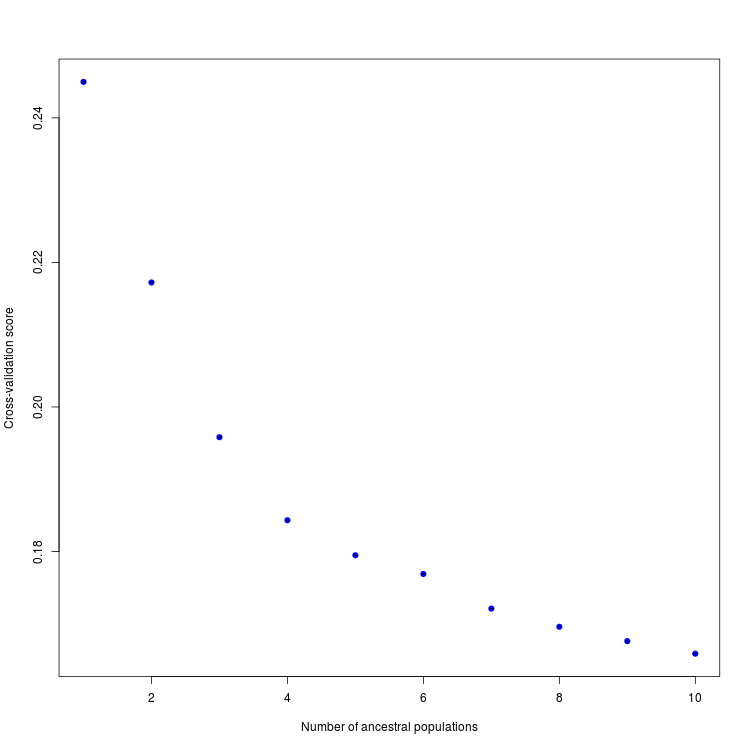

**Fig. S3** A plot of cross-validation scores for increasing numbers of ancestral populations (k) in our TESS3 analyses. This criterion is based on the prediction of a fraction of masked genotypes via matrix completion, which are then compared to masked values considered as the truth. Lower scores are considered as more reliable runs. Scores begin to plateau at k = 4 so we used this value for downstream analyses.

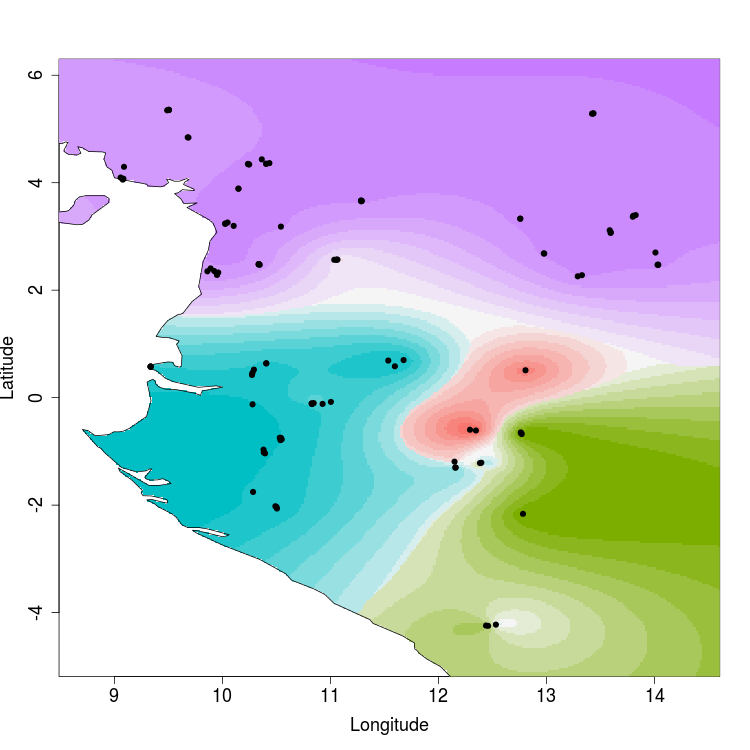

**Fig. S4** A map depicting genetic clusters inferred from the TESS3 analysis when k = 4. This approach takes into account geographic information as well as genetic data. Points represent individuals. Interpolated values of ancestry coefficients are displayed for each of the four clusters inferred and the color gradient corresponds to the level of ancestry.

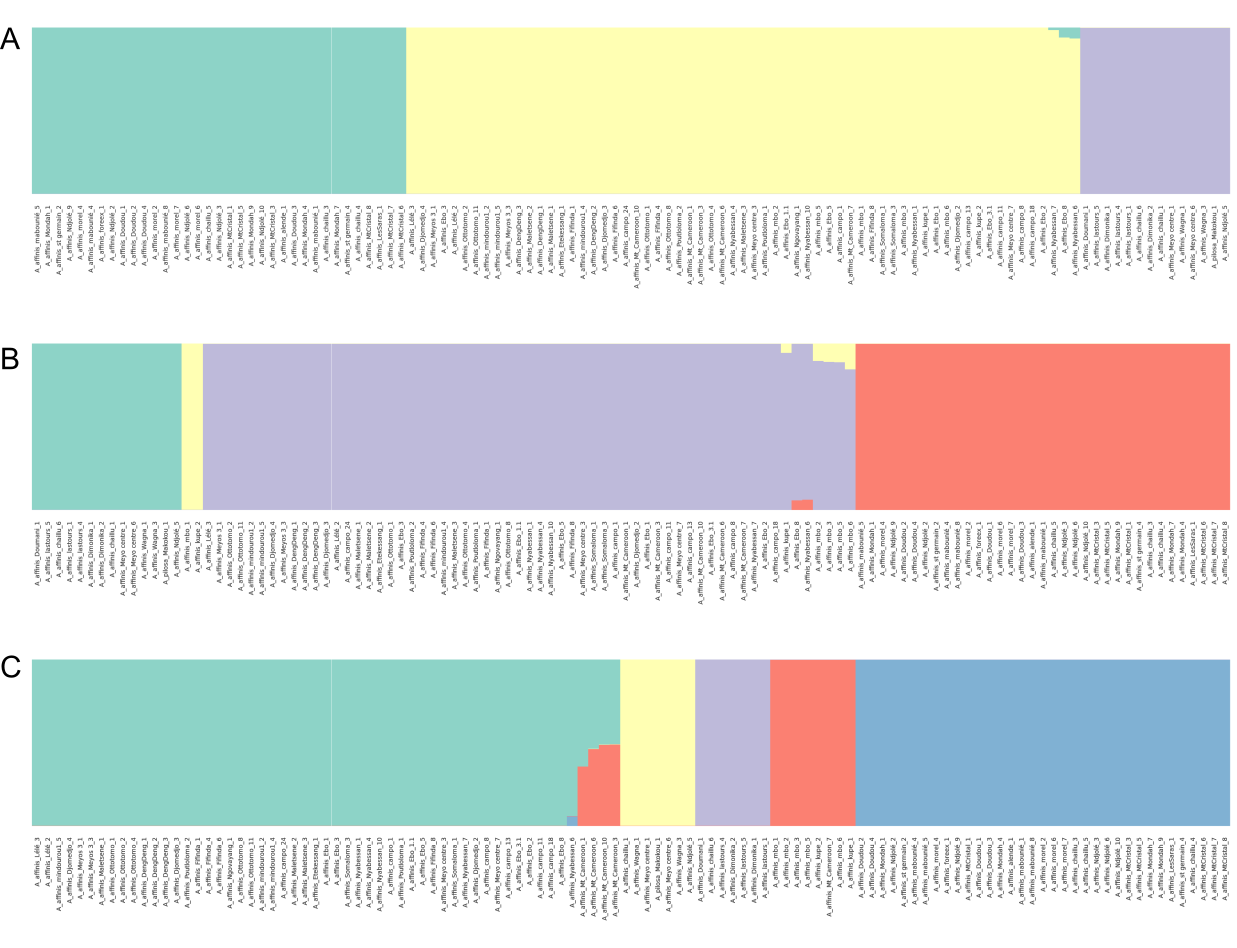

**Fig. S5** Barplots showing the results of Bayesian clustering analyses using fastSTRUCTURE where (A) K = 3, (B) K = 4 and (C) K = 5 based on 257 unlinked SNPs. Three clusters best explained the structure within the dataset while five clusters maximized the marginal likelihood.

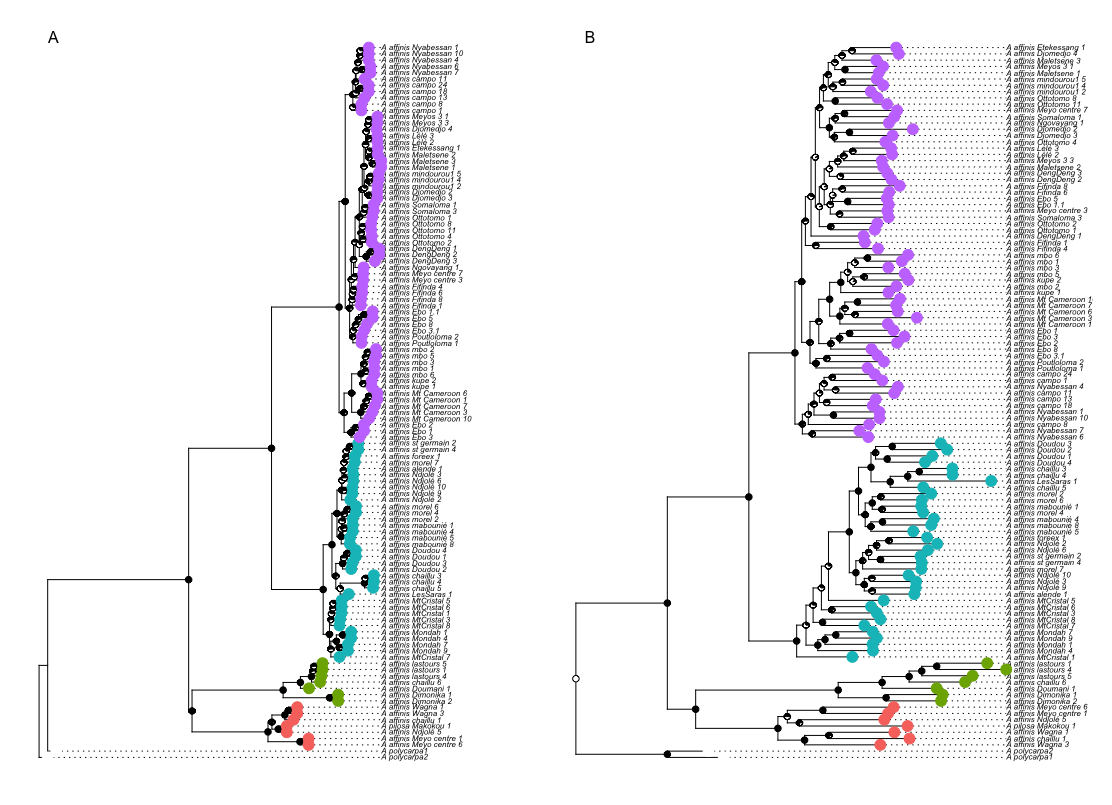

**Fig. S6** Phylogenetic trees of *A. affinis* inferred using the (A) ASTRAL and (B) RAxML. DAPC cluster membership and bootstrap support are shown as colours on tips and as pie charts on nodes respectively. Tip labels for each individual are shown for comparison among trees.

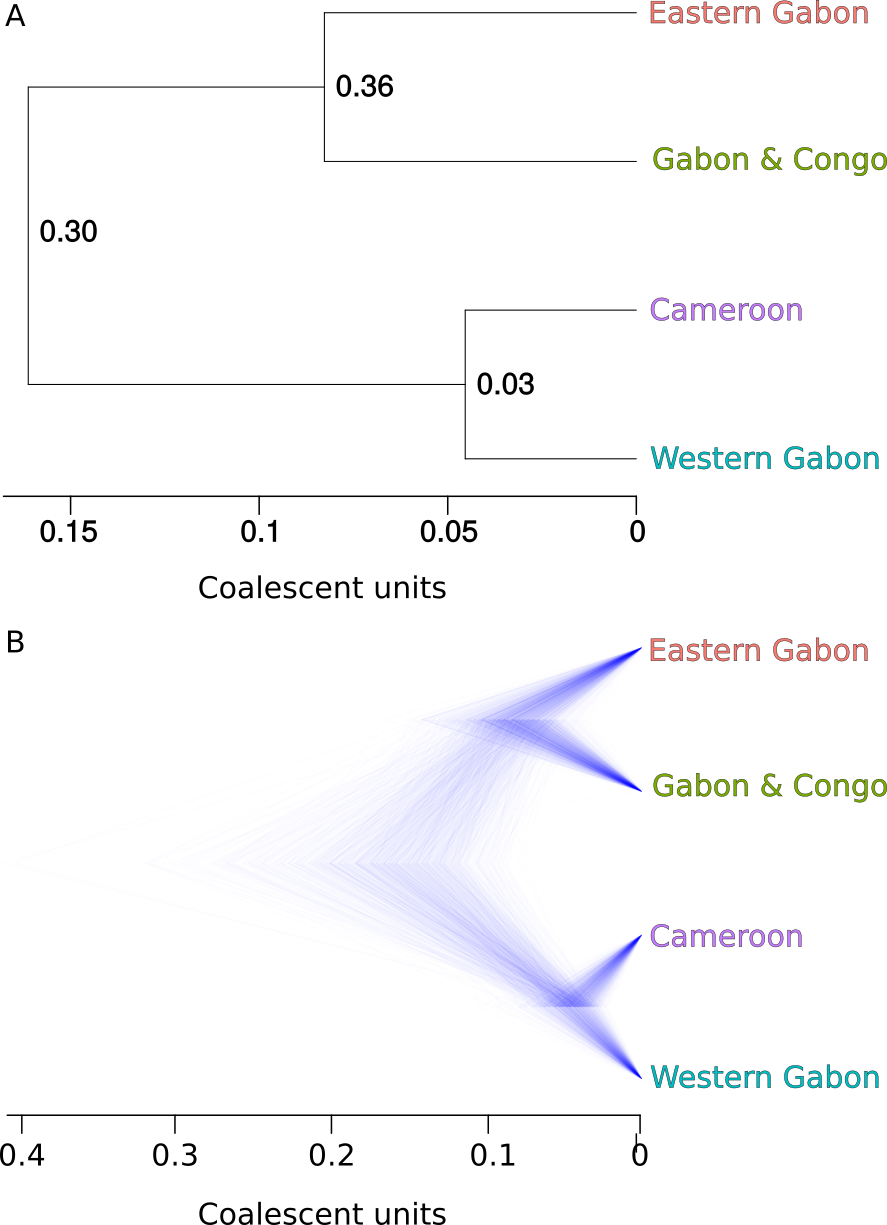

**Figure S7** Phylogenetic tree inferred using the coalescent approach SNAPP with 257 SNPs and 7-10 individuals per DAPC cluster. Panel (A) shows the summary tree with tip labels corresponding to each genetic cluster. Panel (B) shows the posterior distribution of trees of the most common topology inferred by SNAPP.

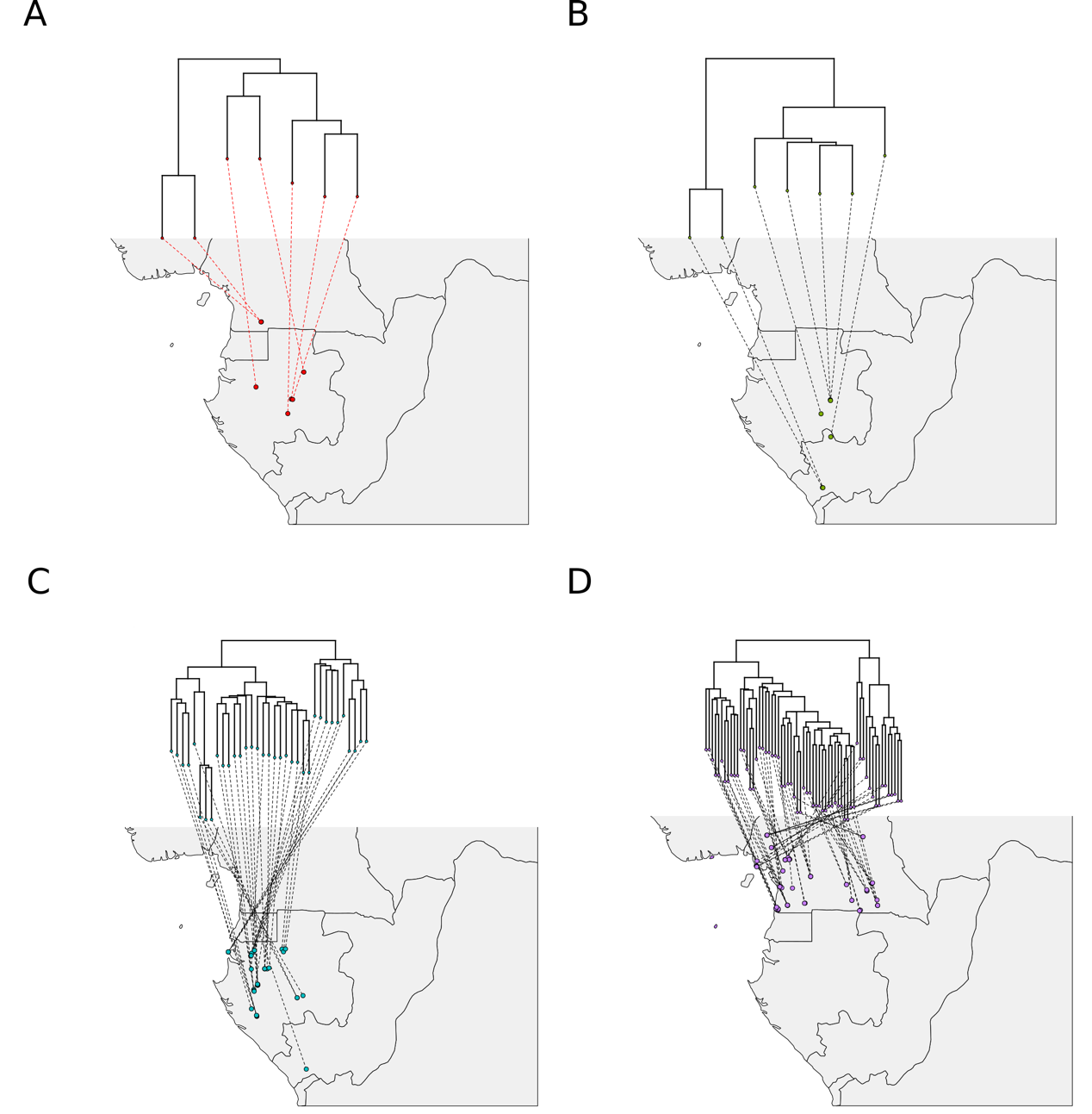

**Fig. S8** Sample locations matched to phylogenetic tree tips. The ASTRAL phylogenetic tree (Fig. 1) was split into clades corresponding to inferred genetic clusters. Circles on the maps indicate locations of individuals in each cluster. Dotted lines connect tips of each tree to their corresponding

location. Clusters 1-4 correspond alphabetically to panes A-D.

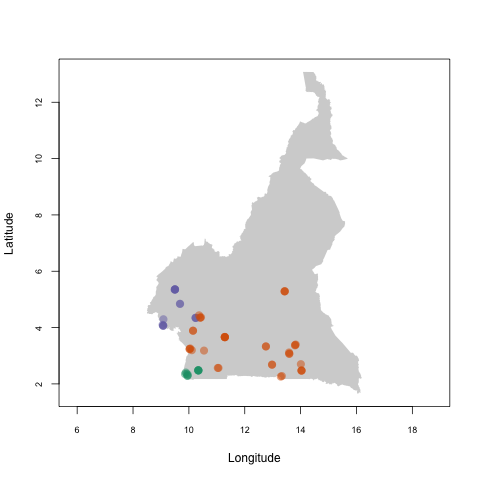
**Fig. S9** DAPC analysis of individuals belonging to Cluster CA shown on a map of Cameroon. We performed the same approach as with the whole dataset and found that k = 3 was the most appropriate number of genetic clusters.

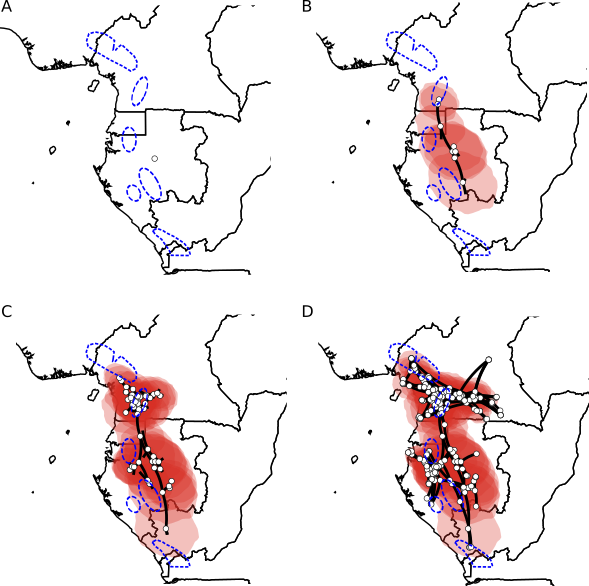
**Fig. S10** Phylogeographic diffusion analysis split into four time slices based on a second set of loci. Images were rendered using spreaD3 and move forward through time starting from the (uncalibrated) time of the most recent common ancestor (A) to the present day (D). White circles represent ancestrally estimated geographic locations for nodes in the inferred phylogenetic tree, as well as current, real locations at tips. Polygons around represent uncertainty of estimated ancestral locations at 80% highest posterior density (HPD). Putative refugia are shown in dashed blue lines.

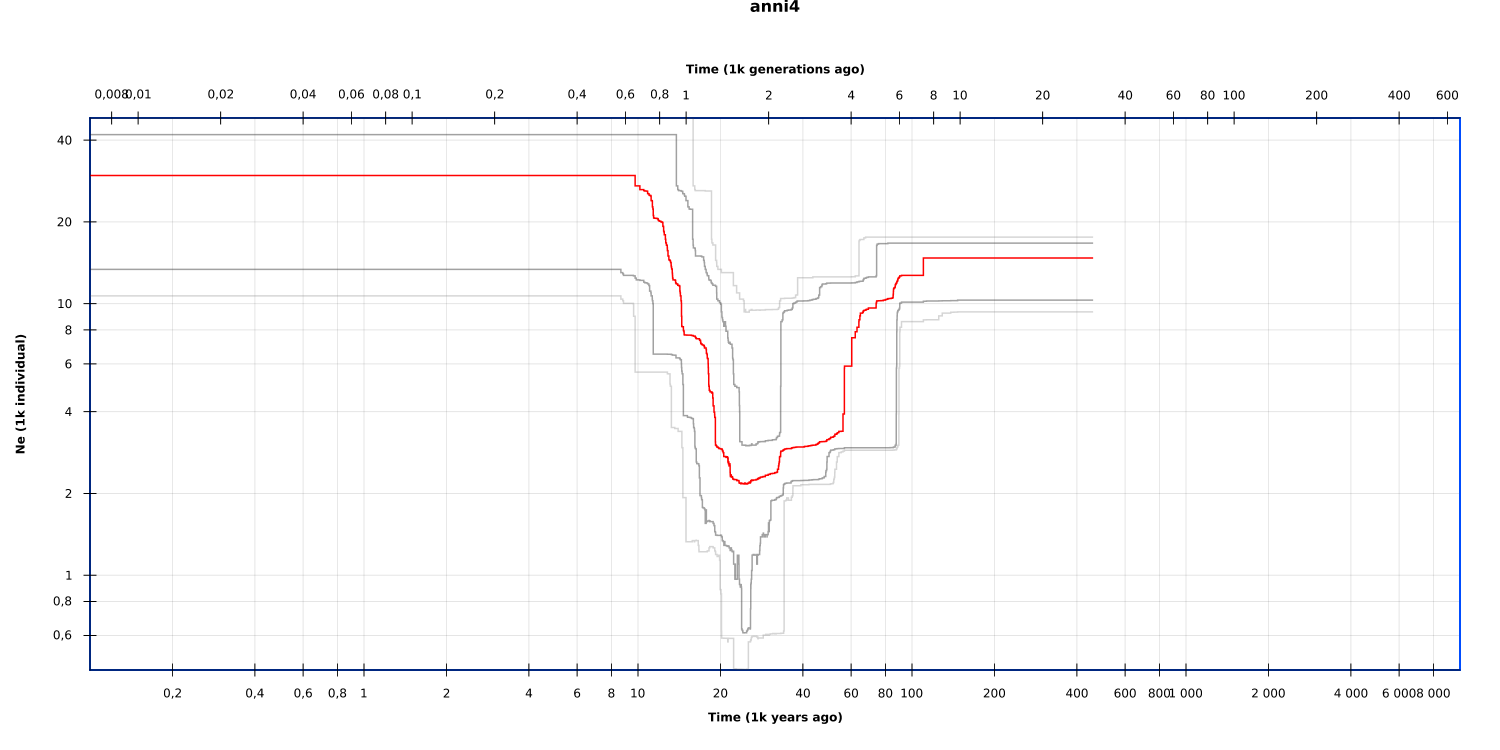

**Figure S11** Stairway plot results for population GC. Red line shows the median *N_e_*. Grey and light grey lines show 80% and 95% confidence intervals respectively.

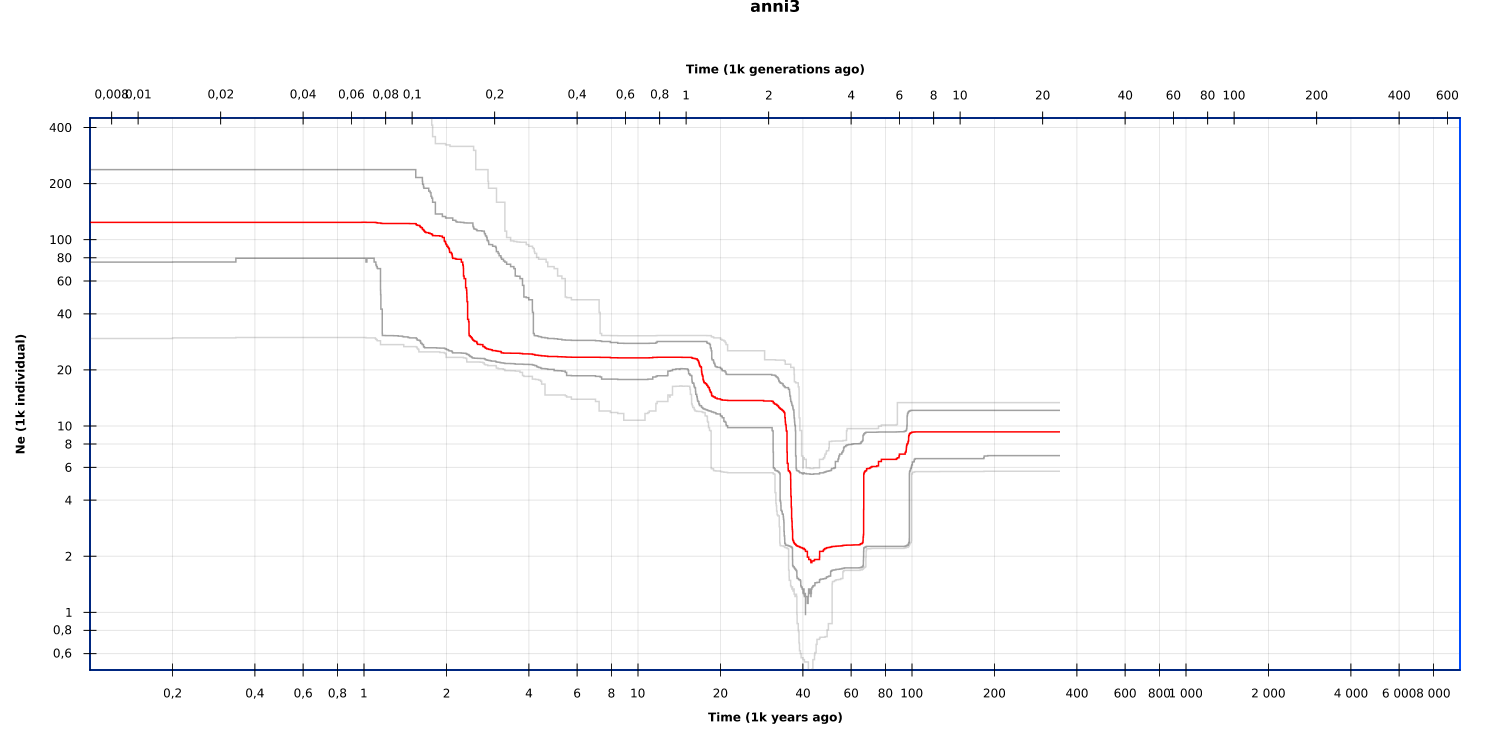

**Figure S12** Stairway plot results for population CA. Red line shows the median *N_e_*. Grey and light grey lines show 80% and 95% confidence intervals respectively.

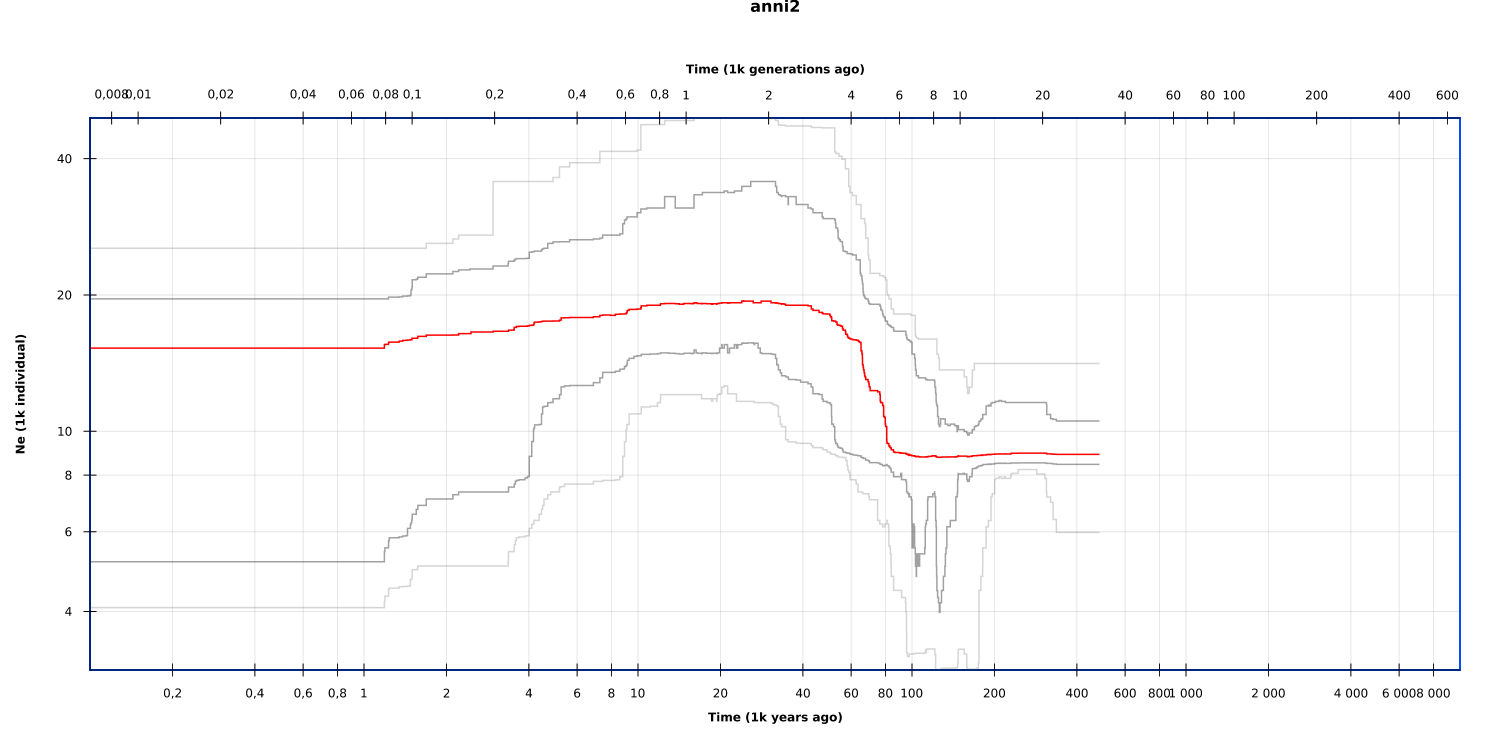
**Figure S13** Stairway plot results for population EG. Red line shows the median *N_e_*. Grey and light grey lines show 80% and 95% confidence intervals respectively.

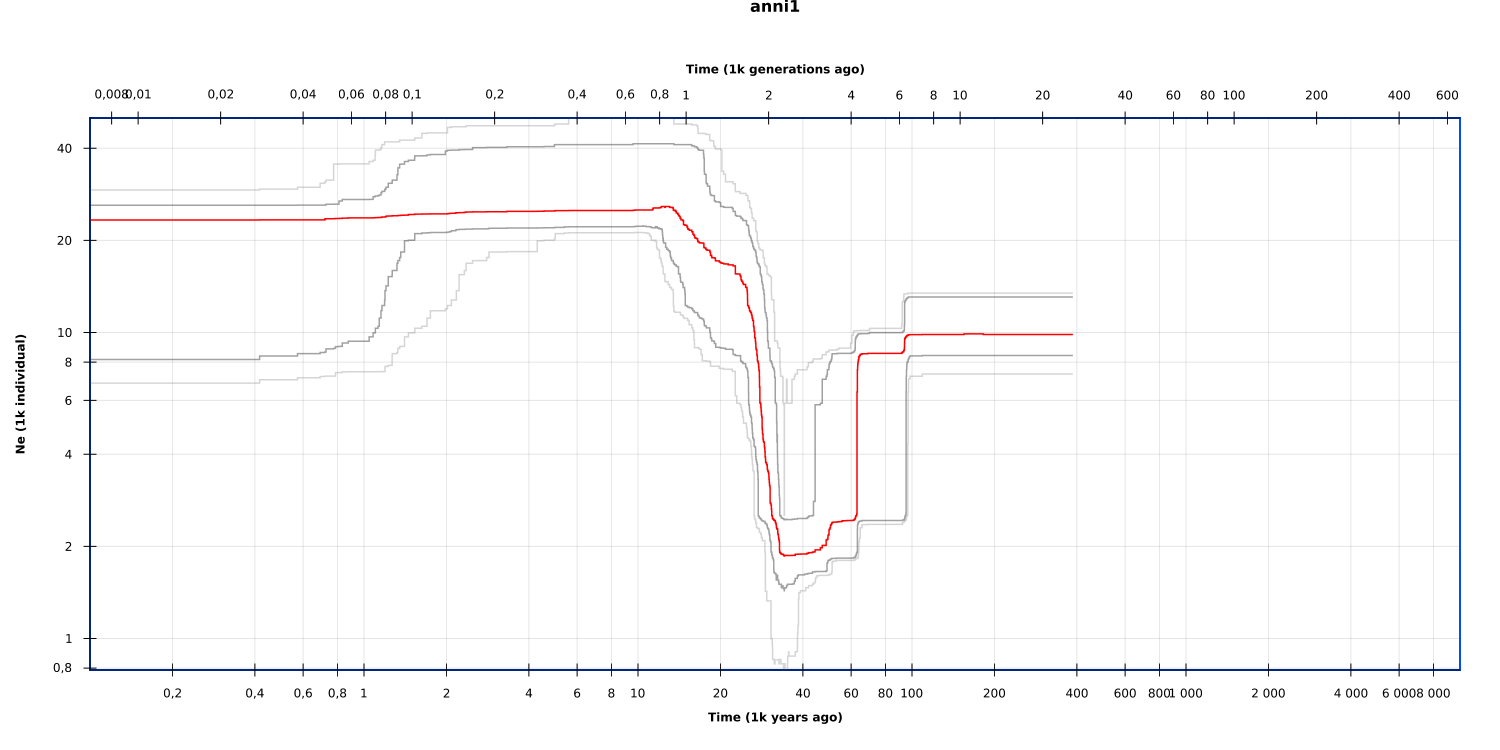
**Figure S14** Stairway plot results for population WG. Red line shows the median *N_e_*. Grey and light grey lines show 80% and 95% confidence intervals respectively.

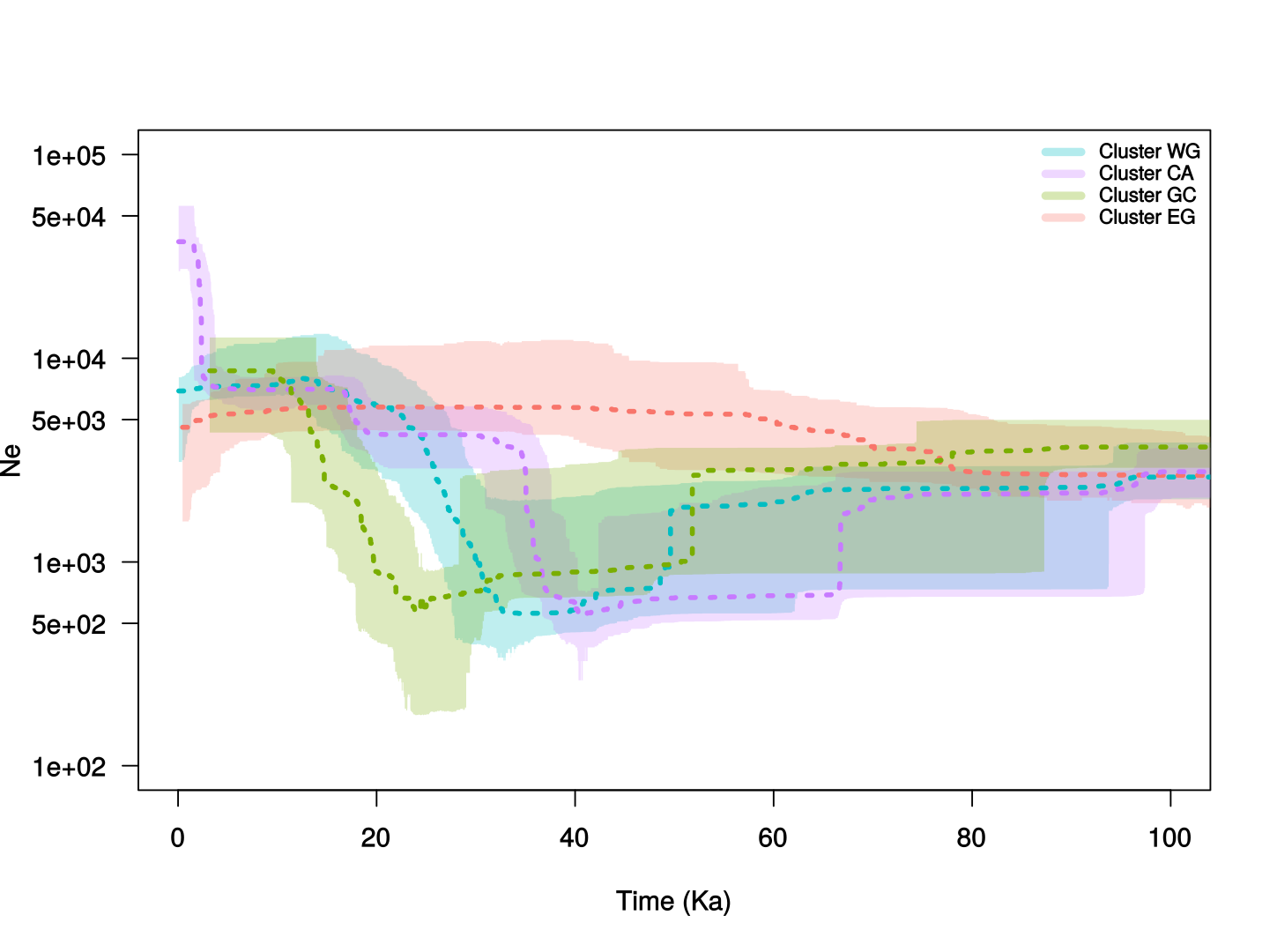

**Figure S15** Stairway plot results for as in figure 3 but using a generation time of 50 years..

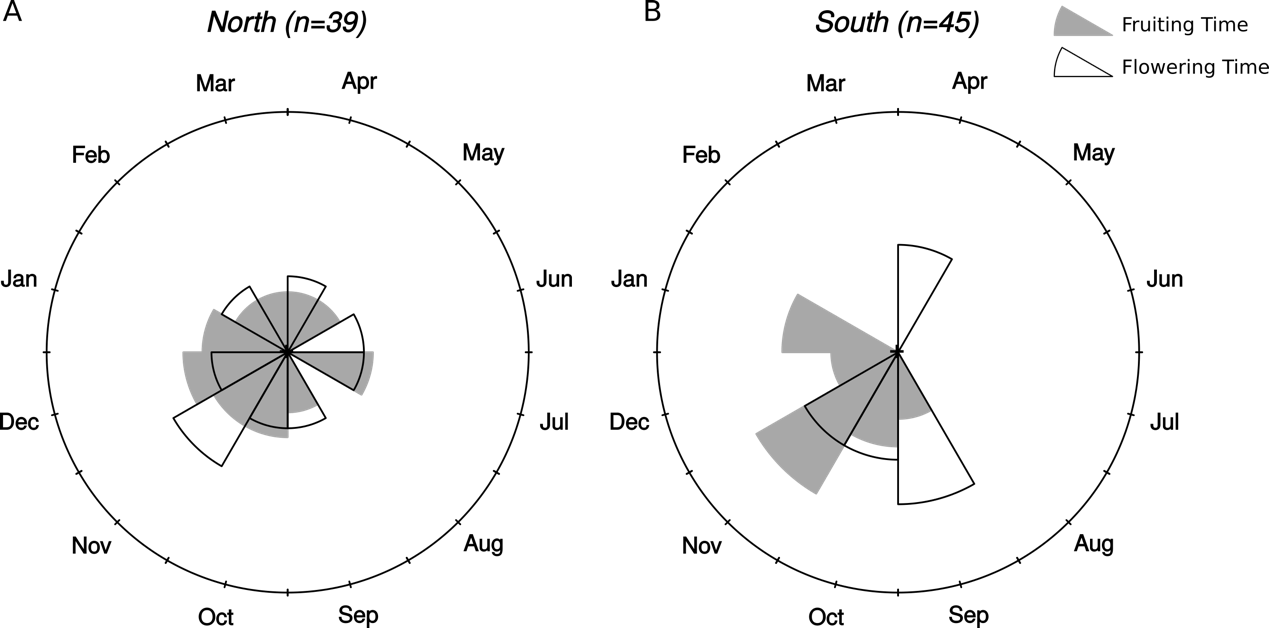

**Figure S16** Plots of months of fruiting and flowering time for *Annickia affinis* individuals found (A) North and (B) South of the climatic inversion. Phenological data was taken from herbarium specimens, where available.

SUPPLEMENTARY TABLES

**Table S1** *Annickia affinis* location data for those specimens used in MaxEnt species distribution modelling (SDM) and those for which DNA was sequenced.

| ID | Tree name | longitude | latitude | SDM | Sequenced | INDEX_TAG |
| --- | --- | --- | --- | --- | --- | --- |
| AA008 | A_affinis_Mt_Cameroon_1 | 9.08051603 | 4.07156001 | x | x | I01_T11 |
| AA010 | A_affinis_Mt_Cameroon_3 | 9.08274704 | 4.07321401 | x | x | I01_T12 |
| AA014 | A_affinis_Ottotomo_1 | 11.288697 | 3.66070803 | x | x | I01_T14 |
| AA015 | A_affinis_Ottotomo_2 | 11.288697 | 3.66070803 | x | x | I01_T15 |
| AA023 | A_affinis_MtCristal_1 | 10.291473 | 0.520887 | x | x | I01_T17 |
| AA025 | A_affinis_Doudou_1 | 10.500082 | -2.033776 | x | x | I01_T18 |
| AA026 | A_affinis_Doudou_2 | 10.504675 | -2.063034 | x | x | I01_T19 |
| AA027 | A_affinis_Doudou_3 | 10.283421 | -1.753833 | x | x | I01_T20 |
| AA028 | A_affinis_Doudou_4 | 10.490086 | -2.02077 | x | x | I01_T21 |
| AA029 | A_affinis_Ebo_1 | 10.235308 | 4.34979804 | x | x | I01_T22 |
| AA030 | A_affinis_Ebo_2 | 10.240975 | 4.34220697 | x | x | I01_T23 |
| AA032 | A_affinis_Mondah_1 | 9.33461904 | 0.57496897 | x | x | I01_T25 |
| AA043 | A_affinis_Somaloma_3 | 12.75694 | 3.331667 | x | x | I01_T29 |
| AA046 | A_affinis_DengDeng_1 | 13.4351629 | 5.29023275 | x | x | I01_T30 |
| AA047 | A_affinis_DengDeng_2 | 13.4316069 | 5.28762501 | x | x | I01_T31 |
| AA048 | A_affinis_DengDeng_3 | 13.4213832 | 5.28367145 | x | x | I01_T32 |
| AA049 | A_affinis_Nyabessan_1 | 10.336509 | 2.48293002 | x | x | I01_T33 |
| AA051 | A_affinis_Nyabessan_4 | 10.336489 | 2.48306899 | x | x | I01_T34 |
| AA053 | A_affinis_Nyabessan_6 | 10.334716 | 2.48508501 | x | x | I01_T35 |
| AA054 | A_affinis_Nyabessan_7 | 10.34537 | 2.47317801 | x | x | I01_T36 |
| AA059 | A_affinis_Djomedjo_2 | 13.596544 | 3.07510867 | x | x | I01_T38 |
| AA060 | A_affinis_Djomedjo_3 | 13.5931982 | 3.06531655 | x | x | I01_T39 |
| AA061 | A_affinis_LesSaras_1 | 12.53181 | -4.2242 | x | x | I01_T40 |
| AA062 | A_affinis_Dimonika_1 | 12.45974 | -4.24668 | x | x | I01_T41 |
| AA065 | A_affinis_st germain_2 | 11.67717 | 0.69962304 | x | x | I01_T43 |
| AA067 | A_affinis_st germain_4 | 11.596572 | 0.58332003 | x | x | I01_T44 |
| AA069 | A_affinis_alende_1 | 11.003973 | -0.078994 | x | x | I01_T45 |
| AA070 | A_affinis_mabounié_1 | 10.548599 | -0.776096 | x | x | I01_T46 |
| AA073 | A_affinis_mabounié_4 | 10.536846 | -0.789789 | x | x | I01_T47 |
| AA074 | A_affinis_mabounié_5 | 10.542062 | -0.748322 | x | x | I01_T48 |
| AA077 | A_affinis_mabounié_8 | 10.530907 | -0.746163 | x | x | I01_T49 |
| AA079 | A_affinis_morel_2 | 10.38052 | -0.967934 | x | x | I01_T50 |
| AA081 | A_affinis_morel_4 | 10.383253 | -1.017661 | x | x | I01_T51 |
| AA083 | A_affinis_morel_6 | 10.383666 | -1.018151 | x | x | I01_T52 |
| AA084 | A_affinis_morel_7 | 10.394331 | -1.036194 | x | x | I01_T53 |
| AA086 | A_affinis_foreex_1 | 11.53526 | 0.69004 | x | x | I01_T54 |
| AA103 | A_affinis_campo_11 | 9.89025 | 2.40468 | x | x | I01_T57 |
| AA105 | A_affinis_Poutloloma_1 | 10.14762 | 3.88916 | x | x | I01_T58 |
| AA106 | A_affinis_Poutloloma_2 | 10.14858 | 3.88842 | x | x | I01_T59 |
| AA114 | A_affinis_Fifinda_8 | 10.04771 | 3.25916 | x | x | I02_T7 |
| AA133 | A_affinis_Meyo centre_3 | 11.05933 | 2.56712 | x | x | I02_T9 |
| AA136 | A_affinis_Meyo centre_6 | 11.06476 | 2.56928 | x | x | I02_T10 |
| AA137 | A_affinis_Meyo centre_7 | 11.03469 | 2.56556 | x | x | I02_T11 |
| AA156 | A_affinis_Ottotomo_8 | 11.285752 | 3.66252498 | x | x | I02_T12 |
| AA159 | A_affinis_Ottotomo_11 | 11.284185 | 3.65316901 | x | x | I02_T13 |
| AA163 | A_affinis_mindourou1_4 | 13.803 | 3.38365233 | x | x | I02_T15 |
| AA164 | A_affinis_mindourou1_5 | 13.8257458 | 3.39717887 | x | x | I02_T16 |
| AA171 | A_affinis_campo_18 | 9.96091667 | 2.32802778 | x | x | I02_T18 |
| AA177 | A_affinis_campo_24 | 9.86061111 | 2.35102778 | x | x | I02_T19 |
| AA179 | A_affinis_Djomedjo_4 | 13.5852222 | 3.11552778 | x | x | I02_T20 |
| AA191 | A_affinis_Meyos 3_1 | 12.97521 | 2.68427 | x | x | I02_T21 |
| AA193 | A_affinis_Meyos 3_3 | 12.97911 | 2.68105 | x | x | I02_T22 |
| AA202 | A_affinis_Maletsene_1 | 14.028 | 2.47273 | x | x | I02_T23 |
| AA203 | A_affinis_Maletsene_2 | 14.03348 | 2.47322 | x | x | I02_T24 |
| AA204 | A_affinis_Maletsene_3 | 14.0345 | 2.47232 | x | x | I02_T25 |
| AA205 | A_affinis_Etekessang_1 | 14.00879 | 2.70069 | x | x | I02_T26 |
| AA257 | A_affinis_mbo_1 | 9.50394304 | 5.35300699 | x | x | I02_T27 |
| AA258 | A_affinis_mbo_2 | 9.50022298 | 5.35422497 | x | x | I02_T28 |
| AA259 | A_affinis_mbo_3 | 9.503535 | 5.35764998 | x | x | I02_T29 |
| AA261 | A_affinis_mbo_5 | 9.50325 | 5.35284 | x | x | I02_T30 |
| AA263 | A_affinis_kupe_1 | 9.68547301 | 4.84066802 | x | x | I02_T32 |
| AA264 | A_affinis_kupe_2 | 9.678424 | 4.84379397 | x | x | I02_T33 |
| AA265 | A_affinis_Mt_Cameroon_7 | 9.08834497 | 4.29533201 | x | x | I02_T34 |
| AA269 | A_affinis_chaillu_1 | 12.14869 | -1.19106 | x | x | I02_T36 |
| AA271 | A_affinis_chaillu_3 | 12.1553 | -1.2999 | x | x | I02_T37 |
| AA273 | A_affinis_chaillu_5 | 12.39647 | -1.2114 | x | x | I02_T39 |
| AA274 | A_affinis_chaillu_6 | 12.38398 | -1.21793 | x | x | I02_T40 |
| AA275 | A_affinis_lastours_1 | 12.77096 | -0.67595 | x | x | I02_T41 |
| AA278 | A_affinis_lastours_4 | 12.76191 | -0.64666 | x | x | I02_T42 |
| AA284 | A_affinis_Ndjolé_3 | 10.82598 | -0.10929 | x | x | I02_T46 |
| AA286 | A_affinis_Ndjolé_5 | 10.84517 | -0.10185 | x | x | I02_T47 |
| AA287 | A_affinis_Ndjolé_6 | 10.82598 | -0.10929 | x | x | I02_T48 |
| AA290 | A_affinis_Ndjolé_9 | 10.92698 | -0.11631 | x | x | I02_T49 |
| AA291 | A_affinis_Ndjolé_10 | 10.27856 | -0.1245 | x | x | I02_T50 |
| AA292 | A_affinis_MtCristal_3 | 10.27856 | 0.46394 | x | x | I02_T51 |
| AA294 | A_affinis_MtCristal_5 | 10.27396 | 0.42789 | x | x | I02_T52 |
| AA295 | A_affinis_MtCristal_6 | 10.27301 | 0.42502 | x | x | I02_T53 |
| AA296 | A_affinis_MtCristal_7 | 10.4055 | 0.6383 | x | x | I02_T54 |
| AA297 | A_affinis_MtCristal_8 | 10.40551 | 0.63833 | x | x | I02_T55 |
| AA300 | A_affinis_Mondah_7 | 9.3351 | 0.57889 | x | x | I02_T56 |
| AA302 | A_affinis_Mondah_9 | 9.3351 | 0.57889 | x | x | I02_T57 |
| AA306 | A_affinis_Ebo_3.1 | 10.405001 | 4.35369504 | x | x | I02_T59 |
| AA308 | A_affinis_Ebo_5 | 10.434563 | 4.36423 | x | x | I02_T60 |
| AA311 | A_affinis_Ebo_8 | 10.3644941 | 4.434456 | x | x | I02_T61 |
| AA312 | A_affinis_Doumani_1 | 12.78239 | -2.16223 | x | x | I02_T62 |
| AA005 |  | 13.289507 | 2.25792403 | x |  |  |
| AA009 |  | 9.07669103 | 4.05760598 | x |  |  |
| AA011 |  | 9.08539798 | 4.07461404 | x |  |  |
| AA012 |  | 9.08604196 | 4.07493398 | x |  |  |
| AA016 |  | 11.288697 | 3.66070803 | x |  |  |
| AA019 |  | 10.7756 | 3.90322 | x |  |  |
| AA020 |  | 10.77278 | 3.90835 | x |  |  |
| AA024 |  | 10.413643 | 0.61811599 | x |  |  |
| AA033 |  | 9.33576099 | 0.57419197 | x |  |  |
| AA034 |  | 9.33618 | 0.57335 | x |  |  |
| AA036 |  | 10.824904 | -0.122868 | x |  |  |
| AA038 |  | 11.699455 | 3.41828001 | x |  |  |
| AA040 |  | 13.56639 | 3.089444 | x |  |  |
| AA042 |  | 12.75944 | 3.328333 | x |  |  |
| AA044 |  | 14.91556 | 3.918611 | x |  |  |
| AA050 |  | 10.342929 | 2.48225503 | x |  |  |
| AA052 |  | 10.341484 | 2.47707903 | x |  |  |
| AA055 |  | 10.34537 | 2.47317801 | x |  |  |
| AA056 |  | 10.34537 | 2.47317801 | x |  |  |
| AA058 |  | 13.5818762 | 3.06229961 | x |  |  |
| AA064 |  | 11.678632 | 0.70037297 | x |  |  |
| AA066 |  | 11.59731 | 0.587068 | x |  |  |
| AA068 |  | 11.595885 | 0.58444899 | x |  |  |
| AA071 |  | 10.547452 | -0.782489 | x |  |  |
| AA072 |  | 10.548483 | -0.78171 | x |  |  |
| AA075 |  | 10.539182 | -0.747032 | x |  |  |
| AA076 |  | 10.5386 | -0.745949 | x |  |  |
| AA078 |  | 10.379351 | -0.967306 | x |  |  |
| AA080 |  | 10.380703 | -0.96708 | x |  |  |
| AA082 |  | 10.38319 | -1.019749 | x |  |  |
| AA085 |  | 11.48695 | 0.60097 | x |  |  |
| AA088 |  | 9.37825 | 4.62247 | x |  |  |
| AA089 |  | 10.14835 | 3.88042 | x |  |  |
| AA090 |  | 10.14888 | 3.88019 | x |  |  |
| AA091 |  | 10.14842 | 3.88578 | x |  |  |
| AA094 |  | 9.94678 | 2.28507 | x |  |  |
| AA096 |  | 9.94928 | 2.28006 | x |  |  |
| AA097 |  | 9.94726 | 2.27985 | x |  |  |
| AA098 |  | 9.94822 | 2.27895 | x |  |  |
| AA099 |  | 9.94824 | 2.29043 | x |  |  |
| AA101 |  | 9.94578 | 2.29207 | x |  |  |
| AA104 |  | 9.88872 | 2.40424 | x |  |  |
| AA108 |  | 10.02546 | 3.2348 | x |  |  |
| AA109 |  | 10.02556 | 3.23881 | x |  |  |
| AA111 |  | 10.10713 | 3.19762 | x |  |  |
| AA113 |  | 10.1036 | 3.1953 | x |  |  |
| AA115 |  | 10.04294 | 3.26029 | x |  |  |
| AA116 |  | 10.0403 | 3.26083 | x |  |  |
| AA117 |  | 10.53042 | 2.78201 | x |  |  |
| AA119 |  | 10.53104 | 2.78108 | x |  |  |
| AA120 |  | 10.53117 | 2.7806 | x |  |  |
| AA121 |  | 10.60579 | 2.8143 | x |  |  |
| AA122 |  | 10.60481 | 2.81539 | x |  |  |
| AA124 |  | 10.60352 | 2.81882 | x |  |  |
| AA125 |  | 10.60599 | 2.82233 | x |  |  |
| AA126 |  | 10.60689 | 2.82368 | x |  |  |
| AA127 |  | 11.12484 | 2.83746 | x |  |  |
| AA128 |  | 11.12479 | 2.83754 | x |  |  |
| AA130 |  | 11.12228 | 2.83495 | x |  |  |
| AA132 |  | 11.05586 | 2.56607 | x |  |  |
| AA134 |  | 11.06009 | 2.56792 | x |  |  |
| AA135 |  | 11.06538 | 2.5696 | x |  |  |
| AA138 |  | 11.35728 | 2.23914 | x |  |  |
| AA139 |  | 11.35735 | 2.23955 | x |  |  |
| AA140 |  | 11.3599 | 2.24558 | x |  |  |
| AA141 |  | 11.35983 | 2.24538 | x |  |  |
| AA142 |  | 11.3598 | 2.24535 | x |  |  |
| AA143 |  | 11.35434 | 2.23501 | x |  |  |
| AA144 |  | 11.43209 | 2.30676 | x |  |  |
| AA145 |  | 11.43181 | 2.3069 | x |  |  |
| AA147 |  | 11.43165 | 2.30848 | x |  |  |
| AA149 |  | 11.4288 | 2.31179 | x |  |  |
| AA150 |  | 11.42969 | 2.3087 | x |  |  |
| AA151 |  | 11.43105 | 2.307 | x |  |  |
| AA152 |  | 11.43633 | 2.30688 | x |  |  |
| AA153 |  | 13.61236 | 3.12561 | x |  |  |
| AA154 |  | 11.284231 | 3.65697197 | x |  |  |
| AA155 |  | 11.286606 | 3.66131203 | x |  |  |
| AA157 |  | 11.285666 | 3.66420102 | x |  |  |
| AA158 |  | 11.284417 | 3.66244401 | x |  |  |
| AA160 |  | 13.7991318 | 3.36641001 | x |  |  |
| AA162 |  | 13.8137449 | 3.38164925 | x |  |  |
| AA165 |  | 13.81978 | 3.38659 | x |  |  |
| AA167 |  | 9.94936111 | 2.28469444 | x |  |  |
| AA168 |  | 9.94858333 | 2.28358333 | x |  |  |
| AA169 |  | 9.94794444 | 2.28166667 | x |  |  |
| AA170 |  | 9.94594444 | 2.28258333 | x |  |  |
| AA172 |  | 9.93677778 | 2.33616667 | x |  |  |
| AA173 |  | 9.89375 | 2.39486111 | x |  |  |
| AA174 |  | 10.0240278 | 2.38283333 | x |  |  |
| AA175 |  | 10.0240278 | 2.39486111 | x |  |  |
| AA176 |  | 9.95011111 | 2.28594444 | x |  |  |
| AA178 |  | 10.2135833 | 2.34177778 | x |  |  |
| AA180 |  | 12.14551 | 2.73426 | x |  |  |
| AA181 |  | 12.1445 | 2.72773 | x |  |  |
| AA182 |  | 12.34999 | 2.46082 | x |  |  |
| AA184 |  | 12.34884 | 2.45789 | x |  |  |
| AA185 |  | 12.34495 | 2.45713 | x |  |  |
| AA186 |  | 12.34772 | 2.45595 | x |  |  |
| AA187 |  | 12.66471 | 2.41651 | x |  |  |
| AA188 |  | 12.67006 | 2.416 | x |  |  |
| AA189 |  | 12.67258 | 2.41947 | x |  |  |
| AA190 |  | 12.67417 | 2.4258 | x |  |  |
| AA192 |  | 12.9762 | 2.683 | x |  |  |
| AA194 |  | 12.98015 | 2.67498 | x |  |  |
| AA195 |  | 12.98005 | 2.67663 | x |  |  |
| AA196 |  | 12.38161 | 2.44593 | x |  |  |
| AA197 |  | 12.36908 | 2.44039 | x |  |  |
| AA198 |  | 12.36509 | 2.43934 | x |  |  |
| AA199 |  | 12.37776 | 2.44589 | x |  |  |
| AA200 |  | 13.77222 | 2.21991 | x |  |  |
| AA201 |  | 13.77079 | 2.229 | x |  |  |
| AA206 |  | 14.0123 | 2.72189 | x |  |  |
| AA207 |  | 14.01322 | 2.70321 | x |  |  |
| AA208 |  | 14.01564 | 2.70581 | x |  |  |
| AA209 |  | 14.01542 | 2.70669 | x |  |  |
| AA210 |  | 14.01676 | 2.70819 | x |  |  |
| AA211 |  | 14.02326 | 2.71386 | x |  |  |
| AA212 |  | 14.0175 | 2.70971 | x |  |  |
| AA213 |  | 14.02069 | 2.71164 | x |  |  |
| AA214 |  | 14.02236 | 2.71271 | x |  |  |
| AA215 |  | 14.02631 | 2.7134 | x |  |  |
| AA216 |  | 13.98987 | 2.68668 | x |  |  |
| AA217 |  | 13.99121 | 2.6871 | x |  |  |
| AA218 |  | 13.99027 | 2.68706 | x |  |  |
| AA219 |  | 13.98814 | 2.69197 | x |  |  |
| AA220 |  | 13.98916 | 2.69386 | x |  |  |
| AA221 |  | 13.89554 | 2.96637 | x |  |  |
| AA222 |  | 13.89681 | 2.96735 | x |  |  |
| AA223 |  | 13.8969 | 2.96896 | x |  |  |
| AA224 |  | 13.88679 | 2.96471 | x |  |  |
| AA225 |  | 13.8991 | 2.99082 | x |  |  |
| AA226 |  | 13.8962 | 2.99049 | x |  |  |
| AA227 |  | 13.89703 | 2.98837 | x |  |  |
| AA228 |  | 13.90792 | 2.98891 | x |  |  |
| AA229 |  | 13.90189 | 2.99286 | x |  |  |
| AA230 |  | 13.90587 | 2.98893 | x |  |  |
| AA231 |  | 13.43303 | 2.43892 | x |  |  |
| AA232 |  | 13.43421 | 2.43854 | x |  |  |
| AA233 |  | 13.43483 | 2.43605 | x |  |  |
| AA234 |  | 13.43444 | 2.43483 | x |  |  |
| AA235 |  | 13.43011 | 2.43499 | x |  |  |
| AA236 |  | 13.42986 | 2.43597 | x |  |  |
| AA237 |  | 13.99604 | 3.27977 | x |  |  |
| AA238 |  | 13.99473 | 3.28112 | x |  |  |
| AA239 |  | 13.99564 | 3.28105 | x |  |  |
| AA240 |  | 13.99646 | 3.28132 | x |  |  |
| AA241 |  | 13.99492 | 3.28218 | x |  |  |
| AA242 |  | 13.99644 | 3.28329 | x |  |  |
| AA243 |  | 13.99726 | 3.28337 | x |  |  |
| AA244 |  | 13.9993 | 3.28274 | x |  |  |
| AA245 |  | 13.33214 | 3.64785 | x |  |  |
| AA246 |  | 13.33898 | 3.64898 | x |  |  |
| AA247 |  | 13.33982 | 3.64959 | x |  |  |
| AA248 |  | 13.34367 | 3.64974 | x |  |  |
| AA249 |  | 13.43539 | 3.64992 | x |  |  |
| AA250 |  | 13.35036 | 3.64891 | x |  |  |
| AA251 |  | 13.10231 | 4.18499 | x |  |  |
| AA252 |  | 13.10395 | 4.18395 | x |  |  |
| AA253 |  | 13.1054 | 4.18379 | x |  |  |
| AA254 |  | 13.10994 | 4.18549 | x |  |  |
| AA255 |  | 13.1101 | 4.18673 | x |  |  |
| AA256 |  | 13.10874 | 4.18674 | x |  |  |
| AA260 |  | 9.48775196 | 5.346867 | x |  |  |
| AA266 |  | 9.04994496 | 4.08558302 | x |  |  |
| AA267 |  | 9.050843 | 4.08845801 | x |  |  |
| AA270 |  | 12.15919 | -1.30014 | x |  |  |
| AA276 |  | 12.76582 | -0.6578 | x |  |  |
| AA277 |  | 12.76544 | -0.66273 | x |  |  |
| AA285 |  | 10.82598 | -0.10929 | x |  |  |
| AA288 |  | 10.82598 | -0.10929 | x |  |  |
| AA289 |  | 10.84517 | -0.1031 | x |  |  |
| AA293 |  | 10.27856 | 0.46394 | x |  |  |
| AA299 |  | 9.3351 | 0.57889 | x |  |  |
| AA301 |  | 9.3351 | 0.57889 | x |  |  |
| AA303 |  | 9.3351 | 0.57889 | x |  |  |
| AA305 |  | 10.404407 | 4.352078 | x |  |  |
| AA307 |  | 10.409601 | 4.34466303 | x |  |  |
| AA309 |  | 10.3887 | 4.39542401 | x |  |  |
| AA310 |  | 10.364493 | 4.434439 | x |  |  |
| Andel, T.R. van 3784 |  | 10.0742 | 2.40833 | x |  |  |
| Bates, G.L. 1959 |  | 11.9 | 3.89333 | x |  |  |
| Bidault, E. 1231 |  | 10.5417 | -0.765278 | x |  |  |
| Bidault, E. 1319 |  | 10.5994 | -0.720278 | x |  |  |
| Bouquet, A. 1921 |  | 12.1428 | -4.10733 | x |  |  |
| Brenan, J.P.M. 8610 |  | 5.25 | 6.33333 | x |  |  |
| Breteler, F.J. 11135 |  | 10.75 | -0.083333 | x |  |  |
| Breteler, F.J. 14597 |  | 9.93333 | -2.7 | x |  |  |
| Briey, J. de 218 |  | 12.85 | -4.86667 | x |  |  |
| Chevalier, A.J.B. 26588 |  | 10.2833 | -0.683333 | x |  |  |
| Couvreur, T.L.P. 400 |  | 10.5416 | 3.18267 | x |  |  |
| Couvreur, T.L.P. 469 |  | 10.1098 | 3.26751 | x |  |  |
| Couvreur, T.L.P. 492 |  | 13.2912 | 2.2608 | x |  |  |
| Couvreur, T.L.P. 519 |  | 9.07668 | 4.05754 | x |  |  |
| Couvreur, T.L.P. 530 |  | 9.33495 | 0.575031 | x |  |  |
| Couvreur, T.L.P. 557 |  | 10.8293 | -0.119966 | x |  |  |
| Couvreur, T.L.P. 591 |  | 11.4674 | 0.422805 | x |  |  |
| Couvreur, T.L.P. 621 |  | 10.2476 | 4.33918 | x |  |  |
| Couvreur, T.L.P. 657 |  | 10.7723 | 3.90717 | x |  |  |
| Couvreur, T.L.P. 671 |  | 10.3367 | 2.47813 | x |  |  |
| Donis, C. 2308 |  | 13.0667 | -5.63333 | x |  |  |
| Etuge, M. 2049 |  | 9.71667 | 4.73333 | x |  |  |
| Etuge, M. 5139 |  | 11.58 | 3.63 | x |  |  |
| Gavage 67 |  | 9.38333 | 0.533333 | x |  |  |
| Gossweiler, J. 6675 |  | 12.5667 | -4.75 | x |  |  |
| Gossweiler, J. 7586 |  | 12.7667 | -4.65 | x |  |  |
| Gossweiler, J. 8085 |  | 12.65 | -4.58333 | x |  |  |
| Hallé, F. 4190 |  | 9.9 | 2.65 | x |  |  |
| Hallé, F. 4201 |  | 9.9 | 2.65 | x |  |  |
| Hédin, L. 1677 |  | 11.5 | 3.51667 | x |  |  |
| Hombert, J. 223 |  | 13.0667 | -5.63333 | x |  |  |
| Hombert, J. 249 |  | 13.0667 | -5.63333 | x |  |  |
| INEF NA |  | 9.38333 | 0.533333 | x |  |  |
| Le Testu, G.M.P.C. 1783 |  | 11 | -2.83333 | x |  |  |
| Leeuwenberg, A.J.M. 7353 |  | 13.6833 | 4.38333 | x |  |  |
| Leeuwenberg, A.J.M. 7355 |  | 13.6833 | 4.38333 | x |  |  |
| Leeuwenberg, A.J.M. 7787 |  | 13.6833 | 4.38333 | x |  |  |
| Letouzey, R. 11751 |  | 13.3 | 2.41667 | x |  |  |
| Letouzey, R. 12361 |  | 10.9667 | 3.81667 | x |  |  |
| Letouzey, R. 5412 |  | 14.8 | 3.16667 | x |  |  |
| Mabiala 790 |  | 12.1 | -4.53333 | x |  |  |
| Mahieu, J. 37 |  | 13.0667 | -5.63333 | x |  |  |
| Mahieu, J. 52 |  | 13.0667 | -5.63333 | x |  |  |
| McPherson, G.D. 13702 |  | 11.5 | -0.416667 | x |  |  |
| McPherson, G.D. 15633 |  | 11.75 | -0.45 | x |  |  |
| Morel, J. 114 |  | 9.75 | 0.25 | x |  |  |
| Namur, C. de NA |  | 12.15 | -4.08333 | x |  |  |
| Nemba, J. 102 |  | 9.6 | 4.91667 | x |  |  |
| Nemba, J. 78 |  | 9.4 | 4.65 | x |  |  |
| Quiroz-Villarreal, D.K. 1725 |  | 11.1803 | -1.06361 | x |  |  |
| Quiroz-Villarreal, D.K. 963 |  | 9.455 | 0.402778 | x |  |  |
| Reitsma, J.M. 1149 |  | 10.5833 | -2.6 | x |  |  |
| Reitsma, J.M. 1723 |  | 10.5667 | -2.58333 | x |  |  |
| Reitsma, J.M. 2205 |  | 11.3667 | 0.733333 | x |  |  |
| Reitsma, J.M. 2347 |  | 11.55 | -0.5 | x |  |  |
| Sargos, R. 132 |  | 12 | -4 | x |  |  |
| Service Forestier du Cameroun 84 |  | 11.3167 | 3.65 | x |  |  |
| Simons, E.L.A.N. 640 |  | 9.48333 | 0.416667 | x |  |  |
| Sosef, M.S.M. 1795 |  | 9.48333 | 0.416667 | x |  |  |
| Sosef, M.S.M. 1881 |  | 11.8733 | 1.08 | x |  |  |
| Sosef, M.S.M. 1906 |  | 11.9267 | 1.29167 | x |  |  |
| Sosef, M.S.M. 1971 |  | 11.785 | 1.36833 | x |  |  |
| Soyaux, H. 125 |  | 9.5 | 0.416667 | x |  |  |
| Stévart, T.O.B.E.B. 4666 |  | 10.5458 | -0.779722 | x |  |  |
| Thomas, D.W. 4345 |  | 8.83333 | 5.01667 | x |  |  |
| Thomas, D.W. 6327 |  | 9.2 | 4.58333 | x |  |  |
| Toussaint, L. 2111 |  | 13.0667 | -5.63333 | x |  |  |
| Toussaint, L. 317 |  | 13.0667 | -5.63333 | x |  |  |
| Towns, A.M. 1099 |  | 12.0694 | 2.18028 | x |  |  |
| Towns, A.M. 743 |  | 9.455 | 0.402778 | x |  |  |
| Towns, A.M. 919 |  | 10.1061 | 0.347778 | x |  |  |
| Towns, A.M. 944 |  | 10.1469 | 0.348056 | x |  |  |
| Wagemans, J. 1867 |  | 13.0667 | -5.63333 | x |  |  |
| Waterman, P.G. 844 |  | 10.0306 | 3.78056 | x |  |  |
| Wieringa, J.J. 5182 |  | 10.8563 | -1.32367 | x |  |  |
| Wilde, J.J.F.E. de 8492 |  | 11.1333 | 2.81667 | x |  |  |
| Wilde, W.J.J.O. de 1499 |  | 10.45 | 3.83333 | x |  |  |
| Wilks, C.M. 2119 |  | 9.38028 | -0.033333 | x |  |  |
| Williamson, E.A. 98 |  | 11.5 | -0.25 | x |  |  |
| Zenker, G.A. 3839 |  | 10.4167 | 3.08333 | x |  |  |
| Zenker, G.A. 441 |  | 10.3833 | 3.06667 | x |  |  |
| AA001 | A_pilosa_Makokou_1 | 12.8056 | 0.5095 |  | x | I01_T7 |
| AA003 | A_affinis_Ngovayang_1 | 10.5416 | 3.1827 |  | x | I01_T8 |
| AA006 | A_affinis_Lélé_2 | 13.328146 | 2.27933904 |  | x | I01_T9 |
| AA007 | A_affinis_Lélé_3 | 13.289402 | 2.25781096 |  | x | I01_T10 |
| AA013 | A_affinis_Mt_Cameroon_6 | 9.0767 | 4.0575 |  | x | I01_T13 |
| AA017 | A_affinis_Ottotomo_4 | 11.28275 | 3.66934 |  | x | I01_T16 |
| AA031 | A_affinis_Ebo_3 | 10.247542 | 4.33912997 |  | x | I01_T24 |
| AA035 | A_affinis_Mondah_4 | 9.335 | 0.575 |  | x | I01_T26 |
| AA037 | A_affinis_Ndjolé_2 | 10.8293 | -0.12 |  | x | I01_T27 |
| AA041 | A_affinis_Somaloma_1 | 12.7575 | 3.329167 |  | x | I01_T28 |
| AA057 | A_affinis_Nyabessan_10 | 10.33666 | 2.47813 |  | x | I01_T37 |
| AA063 | A_affinis_Dimonika_2 | 12.44005 | -4.24089 |  | x | I01_T42 |
| AA093 | A_affinis_campo_1 | 9.94969 | 2.28528 |  | x | I01_T55 |
| AA100 | A_affinis_campo_8 | 9.94824 | 2.29043 |  | x | I01_T56 |
| AA107 | A_affinis_Fifinda_1 | 10.02569 | 3.23364 |  | x | I01_T60 |
| AA110 | A_affinis_Fifinda_4 | 10.02484 | 3.24031 |  | x | I01_T61 |
| AA112 | A_affinis_Fifinda_6 | 10.10397 | 3.19647 |  | x | I01_T62 |
| AA131 | A_affinis_Meyo centre_1 | 11.05205 | 2.56583 |  | x | I02_T8 |
| AA161 | A_affinis_mindourou1_2 | 13.7991318 | 3.36641001 |  | x | I02_T14 |
| AA166 | A_affinis_campo_13 | 9.92358333 | 2.35741667 |  | x | I02_T17 |
| AA262 | A_affinis_mbo_6 | 9.48775 | 5.34687 |  | x | I02_T31 |
| AA268 | A_affinis_Mt_Cameroon_10 | 9.0567 | 4.09647 |  | x | I02_T35 |
| AA272 | A_affinis_chaillu_4 | 12.15947 | -1.30085 |  | x | I02_T38 |
| AA279 | A_affinis_lastours_5 | 12.76544 | -0.66279 |  | x | I02_T43 |
| AA280 | A_affinis_Wagna_1 | 12.34676 | -0.61078 |  | x | I02_T44 |
| AA282 | A_affinis_Wagna_3 | 12.29144 | -0.59678 |  | x | I02_T45 |
| AA304 | A_affinis_Ebo_1.1 | 10.404585 | 4.34993802 |  | x | I02_T58 |

**Table S2** Statistics for datasets recovered by the two pipelines, HybPiper and SeCaPr, used in this study. Phylogenetic informative sites (PIS) and SNPs were only calculated for a single pipeline based on their usage (phylogenetics vs population genetics).

| Pipeline | No. markers | Total length (kb) | Avg length (kb) | No. PIS | No. SNPs |
| --- | --- | --- | --- | --- | --- |
| Hybpiper | 351 | 754.422 | 2.149 | 33787 | N/A |
| SeCaPr | 306 | 181.246 | 0.592 | N/A | 7020 |

**Table S3** Maxent variable contributions as returned by the *variables_importance* function in ‘biomod2’. The closer the values are to 1.00 the greater the influence the variable has on the model.

| Bioclimatic variable | Variable importance |
| --- | --- |
| Annual Mean Temperature (Bio1) | 0.174 |
| Mean Temperature of Warmest Quarter (Bio10) | 0 |
| Mean Temperature of Coldest Quarter (Bio11) | 0.105 |
| Annual Precipitation (Bio12) | 0.475 |
| Precipitation of Wettest Month (Bio13) | 0.909 |
| Precipitation of Driest Month (Bio14) | 0.068 |
| Precipitation Seasonality (Bio15) | 0.534 |
| Precipitation of Wettest Quarter (Bio16) | 0.175 |

**Table S4** Table detailing phenological information on flowering and fruiting time taken from herbarium specimens of *Annickia affinis*.

| North/South | Collector | Month Collected | flowering | fruiting | country | Latitude | Longitude |
| --- | --- | --- | --- | --- | --- | --- | --- |
| South | Gossweiler, J. | September | Y | N | Angola | -4.75 | 12.56666667 |
| South | Gossweiler, J. | October | Y | N | Angola | -4.583333333 | 12.65 |
| South | Le Testu, G.M.P.C. | September | Y | N | Gabon | -2.833333333 | 11 |
| South | Williamson, E.A. |  | Y | Y | Gabon | -0.25 | 11.5 |
| South | Couvreur, T.L.P. | November | y | y | Gabon | -0.119966 | 10.829342 |
| South | Breteler, F.J. | April | Y | N | Gabon | -0.083333333 | 10.75 |
| North | Foury, P. |  | Y | N | Cameroon | 0 | 0 |
| North | Sosef, M.S.M. | February | Y | Y | Gabon | 1.368333333 | 11.785 |
| North | Andel, T.R. van | June | Y | N | Cameroon | 2.4 | 10.06666667 |
| North | Hallé, F. | November | Y | Y | Cameroon | 2.65 | 9.9 |
| North | Hallé, F. | November | Y | Y | Cameroon | 2.65 | 9.9 |
| North | Zenker, G.A. | November | Y | N | Cameroon | 3.066666667 | 10.38333333 |
| North | Letouzey, R.G. | July | Y | Y | Cameroon | 3.166666667 | 14.8 |
| North | Couvreur, T.L.P. | September | y | y | Cameroon | 3.267508 | 10.10982 |
| North | Bates, G.L. |  | Y | Y | Cameroon | 3.893333333 | 11.9 |
| North | Couvreur, T.L.P. | October | y | y | Cameroon | 4.057537 | 9.076678 |
| North | Couvreur, T.L.P. | April | Y | Y | Cameroon | 4.296833 | 9.078546 |
| North | Brenan, J.P.M. | December | Y | N | Nigeria | 6.333333333 | 5.25 |
